# Lymphatic drainage of cerebrospinal fluid using lymph node seeker ^64^Cu-labeled Gram-negative bacterial extracellular vesicles PET

**DOI:** 10.1101/2024.12.04.626900

**Authors:** Kyuwan Kim, Changjin Lee, Ran Ji Yoo, Yoori Choi, Chanwoo Lee, Jaewook Lee, Minseok Suh, Yun-Sang Lee, Yong Song Gho, Dong Soo Lee

## Abstract

Cerebrospinal fluid (CSF) is drained into the systemic lymphatics via paravertebral lymph nodes. Superficial and deep cervical lymph nodes collect CSF in mice, but the exact and quantified routes are unknown. Recently, we simultaneously visualized cervical, sacral and iliac lymph nodes via serial imaging on the intrathecal [^64^Cu]Cu-albumin positron emission tomography. Paravertebral lymph nodes might act as sentinels to monitor the CSF, brain, and spinal cord. We used ^64^Cu-labeled *Escherichia coli* extracellular vesicles, outer membrane vesicles (OMVs), as lymph node seekers for intrathecal administration with an optimized volume/rate and quantified the differential amounts of various paravertebral lymph nodes along the axis of the brain and spinal cord in mice. The quantified results revealed 77.3% in superficial and deep cervical lymph nodes, 11.4% in abdominal/pelvic lymph nodes and 11.3% in sacral lymph nodes. Click-labeled [^64^Cu]Cu-OMVs were drained to reach and stop at the lymph nodes on serial quantification. The cervical lymph nodes drained most of the OMV-laden CSF, which is proportional to the surface areas of the brain (70%) and spinal cord in mice. We propose that all paravertebral lymph nodes monitor the segmental regions of the brain and spinal cord as immediate sentinel lymph nodes against the central nervous system.

## Introduction

Brain waste is drained via cerebrospinal fluid (CSF) to the lymphatics of the body, where it finally reaches the systemic circulation (1–10). Paravertebral lymph nodes are known to be the first stops of the brain-derived fluids and their solutes, which range from small molecules in the CSF to rare macromolecules, even extracellular vesicles (EVs) or cells. The pathway from the CSF to the meningeal lymphatics had been unknown until the dural lymphatics were discovered in humans and mice (4). Since then, investigators have revealed that the barrier layers of arachnoids are not perfect but have defects or intentional tolls (11–13). The first report of defects in the arachnoid barrier cell layer was from a pig study using classical histopathology methods (11). The authors named them cranial arachnoid granulation (CAG)-like dural gaps, which sometimes appear similar to fissures; they likened CAG-like gaps as arachnoid granulations that penetrate into the sagittal sinuses of the dura. As the defects were not expected to exist in the arachnoid barrier cells, no one has previously sought defects in the barrier cell layer.

The necessity of finding drainage routes from the CSF space to the dural meningeal lymphatics encouraged the investigation of defects in the arachnoid barrier cell layer containing claudin-11. While reporting subarachnoid lymphatics-like membrane in mice, Mollgard et al. (12) also reported that there were defects in the arachnoid barrier cell layer, whose continuity is lost in the proximity of dural veins. This finding was corroborated by another report by Smyth et al. (13), where the authors called the arachnoid barrier defects “arachnoid cuff exit” in mice. They revealed the opening of the arachnoid barrier around bridging veins that go outwards from the brain to the dural venous sinuses. MRI contrasts were also found to be dense on the dura side of the bridging veins once these contrast dyes were injected intrathecally into humans. Although the arachnoid cuff exit tends not to allow dura to the subarachnoid entry of fluid/molecules, this structure can be used by the drainage routes of fluid/contrast materials in the cranium.

CSF lymphatic drainage was repeatedly well visualized using various contrast materials of varying sizes and characteristics (3,4,6,10,14). But, the routes or passages via which CSF and molecules reach the exterior of the brain and spinal cord had been unknown. In this context, the discovery of sacral lymph nodes by Kwon et al. (7) and Ma et al. (6) implied diverse routes of lymphatic drainage of the brain and spinal codes after intrathecal (i.t.) and intra-cisterna magna (i.c.m.) injection, respectively. In addition, recent delineation of the passage from cribriform plates to cervical lymph nodes via the nasopharyngeal lymphatic plexus dramatically enhanced our understanding of the routes from the area just outside of the dura to the local lymph nodes (15). However, there is little information about the quantitative ratio of CSF and its contents, intrinsic or exogeneous, along the brain and spinal cord, i.e., whether there are major outlets from the CSF space or drainage from all over the dura contacts of CSF in a distributed manner. To date, there is no evidence of a major or main outlet. Moreover, the drainage characteristics of varying solutes ranging from electrolytes, small molecules, macromolecules, and EVs to cells have not been elucidated.

Previously, we reported that radio-labeled albumin remained in various lymph nodes after an intrathecal administration with the optimized amount and speed of injection in mice (10,16). Considering the very small amount of total CSF and production rate in mice, i.c.m. injection cannot be easily adopted for successful injection and observation. Intrathecal injection should also be adjusted not to remain in the injection site (too small amount and/or too low rate of injection) and not to infuse too much or too fast (16). In repeated experiments, we found the best range of reproducibility for the i.t. method. To have overcome these obstacles, radio-labeled albumin posed another problem, because albumin starts to flow out of the lymph nodes once it reaches them. The problem may have been solved by using the ratio of current lymph node activity over the tentative input function or adopting 24-hour delayed imaging (10).

In this study, we used *Escherichia coli*-derived EVs, i.e., the outer membrane vesicles (OMVs) (17,18), to quantify the lymphatic drainage. OMVs have several advantages, since they are larger particulate materials than albumin, surveil *in vivo*, and are caught in lymph nodes. The OMVs were labeled via click chemistry with dibenzocyclooctyne (DBCO) and chelator-N_3_-radionuclide, so separation was not necessary after mixing the DBCO-labeled OMVs and [^64^Cu]Cu-chelator-N_3_ (19,20). Micro positron emission tomography (microPET) was used to repeatedly acquire whole-body imaging with no background radioactivity (21,22). No radioactive background was obvious we had introduced the radio-labeled tracer ([^64^Cu]Cu-OMVs) to a closed compartment (CSF space). We also demonstrated that unlike OMVs, extracellular vesicle-mimetic nanovesicles (ENVs) (23), which are produced from mammalian cells and labeled in a similar manner to OMVs, can also be used to quantify the lymphatic drainage of CSF but not the lymph node uptake. All activities after administration could be easily read by investigators, and the tissue activity were quantified by PET and their established software.

## Results

### OMV labeling with ^64^Cu

Purified OMVs derived from the *E. coli* BL21(DE3) *ΔmsbB* mutant (18) were radio-labeled to trace their *in vivo* biodistribution across different administration routes. The surfaces of the vesicles were covalently coupled with DBCO molecules to the amine residues of vesicular membrane proteins exposed to the environment. The DBCO-labeled OMVs were subsequently coupled with chelated ^64^Cu radionuclide-NOTA-N_3_ complexes using alkyne-azide click chemistry (Fig. 1A). The radio-labeling efficiency was determined using radio instant thin layer chromatography-silica gel (iTLC-SG) with 0.1 M citric acid as the mobile phase after every radio-labeling procedure. The retention factor (*R*_f_) of ^64^Cu was 0.9-1.0, and that of [^64^Cu]Cu-NOTA-N_3_ was 0.5-0.6. *R*_f_ of ^64^Cu-labeled OMVs was 0.1−0.2, and the radio-labeling efficiency of these vesicles was 99% (Fig. 1B). Thin-layer chromatographic analysis confirmed the preferential chelation of [[Cu with NOTA-N[ and spatial localization of radioactivity on OMVs, which implies the efficient coupling process of alkyne-azide click chemistry and the effective removal of excess free DBCO molecules via diafiltration after the coupling with DBCO on the surface of OMVs.

**Figure 1.**
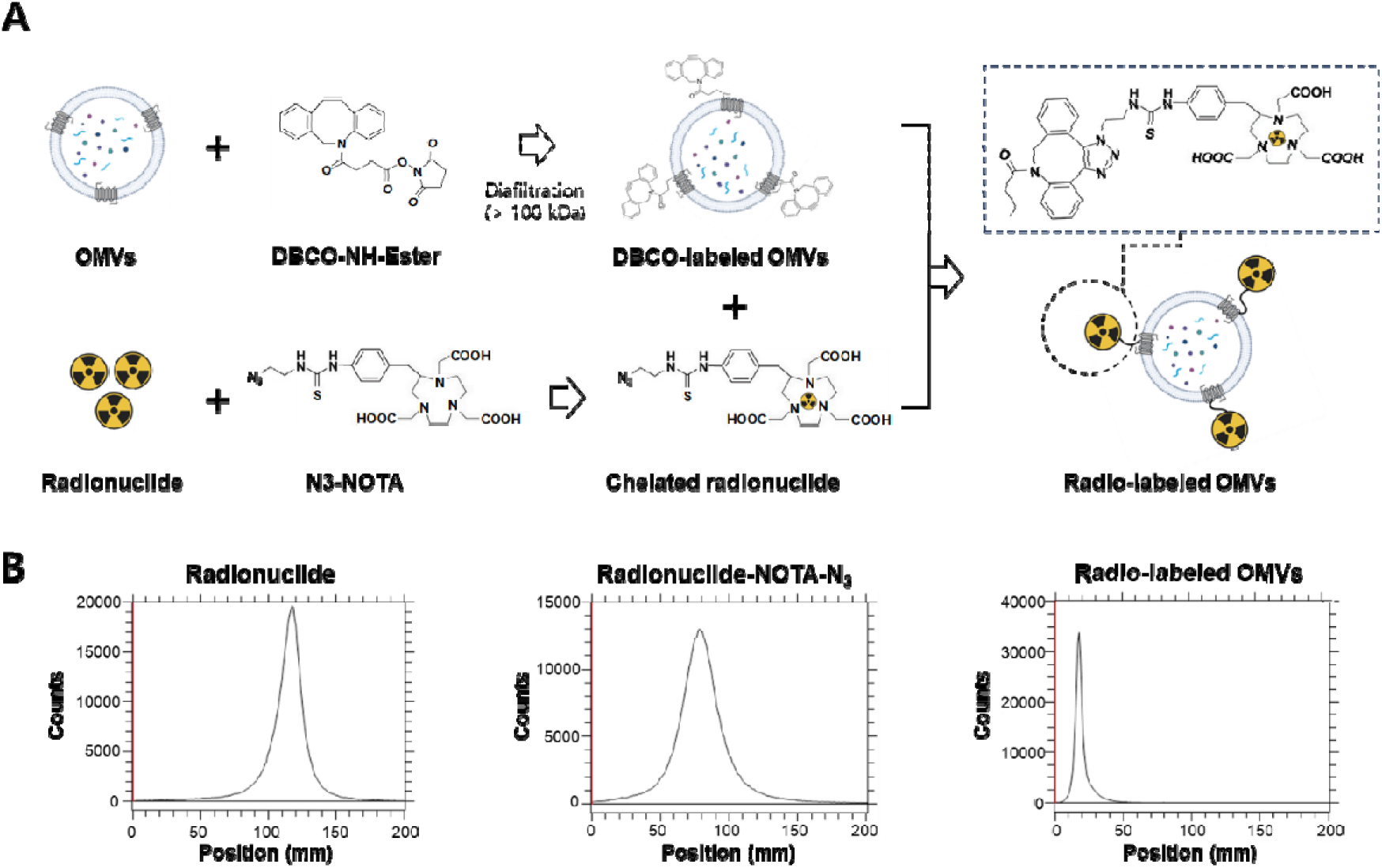
Radio-labeling of OMVs. **(A)** Schematic of radio-labeling on OMVs. Alkyne-tethered OMVs were conjugated with [^64^Cu]Cu-NOTA-N_3_ via click chemistry. **(B)** Labeling efficiency and quality of radio-labeled OMVs were analyzed using instant thin-layer chromatography.

### ^64^Cu-labeled OMVs in the gastrointestinal tract after oral administration

After oral administration, ^64^Cu-labeled OMVs were traced in the gastrointestinal tract (Suppl. Fig. 1, Suppl. Movie 1). From the esophagus to the rectum, the ^64^Cu radioactivity remained in or around the luminal boundary and did not cross the mucosa-epithelial barrier in mice. Beyond the expected half-life of gastric emptying, ^64^Cu radioactivity remained longer. Considering that the mice were fed *ad libitum* between imaging sessions (especially at 2, 4, and 8 hours and the next day), the ^64^Cu radioactivity was late in excretion from the gastric lumen. These results suggest that the ^64^Cu radioactivity remained stable with the OMVs in the gastrointestinal milieu, although the labeled ^64^Cu could be released or recycled in the gastric lumen until 24 or 48 hours after oral administration.

### Differences in the distribution of ^64^Cu-labeled OMVs after subcutaneous and intramuscular administration

After the subcutaneous administration to the right forelimb and right hindlimb, subcutaneous collection activity was maintained around the administration sites until 48 hours (Suppl. Fig. 2A, Suppl. Movie 2A). A very slow flow directed the radioactivity to a dermal flow (or backflow) with scattered localization near and far from the administration site. Since we did not separate free [^64^Cu]Cu-NOTA-N_3_ from ^64^Cu-labeled OMVs, the radioactivity of this rare impurity ([^64^Cu]Cu-NOTA-N_3_) was released via the bladder. Until 8 hours, [^64^Cu]Cu-NOTA-N_3_ was slowly released, which indicates that the lymph flow from the subcutaneous areas after the administration of [^64^Cu]Cu-NOTA-N_3_ took much longer than that after the intravenous administration (Fig. 2A). Notably, only subcutaneous administration via the right hindlimb resulted in signals from the thoracic duct outlet to the subclavian vein. After entering systemic blood circulation, hepatic excretion may result in signals in the intestine and even in the rectum at 8 hours. The remaining subcutaneous radioactivity at the sites of administration suggested that the radioactivity *in situ* was integrated within professional phagocytes (e.g., residential macrophages) or possibly non-professional phagocytes such as stromal and parenchyma cells under the skin.

**Figure 2.**
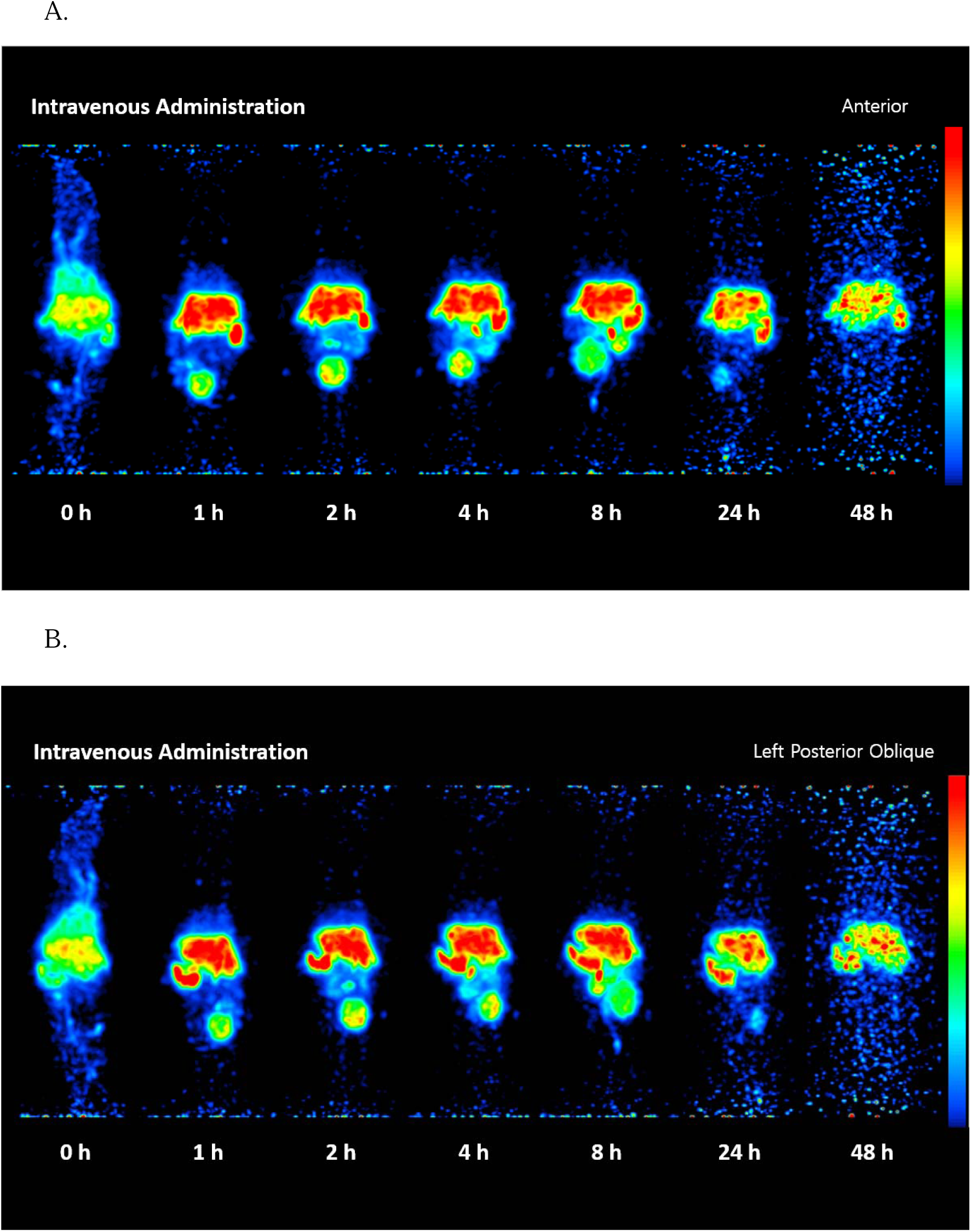
Biodistribution of ^64^Cu-labeled OMVs after intravenous administration in a mouse in the A) anterior and B) left posterior oblique views. The spleen uptake was associated with the liver and persisted throughout the sequential imaging. Small amounts of metabolized [^64^Cu]Cu fragments were excreted via the bladder and intestines (faint abdominal/rectum uptake, C57BL/6, Male, 14 weeks).

Intramuscular administration disclosed a slightly different distribution from that of subcutaneous administration (Suppl. Fig. 2B, Suppl. Movie 2B). The retention of radioactivity at the administration site was identical to that after the subcutaneous administration. The radioactivity appeared in the bladder at 1 hour but rapidly disappeared afterwards (no activity after 2 hours), which suggests that the interstitial [^64^Cu]Cu-NOTA-N_3_ was removed within 2 hours. This initial return of ^64^Cu-labeled OMVs to the systemic blood circulation enabled the ^64^Cu-labeled OMVs to be excreted into the intestine, which resulted in a faint intestinal radioactivity signal. During this normal drainage and excretion of [^64^Cu]Cu-NOTA-N_3_- and ^64^Cu-labeled OMVs, the radioactivity was retained in the lymph node, probably due to the capture of ^64^Cu-labeled OMVs by sentinel lymph nodes. The nearest and brightest lymph node was definitely the regional sentinel, and the second or third might be sequential or simultaneous drainage from the first lymph node.

### Capture of distributed ^64^Cu-labeled OMVs along the communicating lymphatic channels after an intraperitoneal administration

After an intraperitoneal administration, ^64^Cu-labeled OMVs remained in the peritoneum until 48 hours, and the impurities [^64^Cu]Cu-NOTA-N_3_, which were drained from the peritoneum to the systemic blood circulation, were excreted via the kidneys and bladder within 2 hours (Suppl. Fig. 3, Suppl. Movie 3). From the beginning, ^64^Cu-labeled OMVs drained from the peritoneum to the pleural cavity via diaphragmatic lacunae and stomata (that connects the pleural cavity and peritoneum) and were captured by scattered lymph nodes around the pleural cavity and mediastinum that reached the cervical lymph nodes. The radioactivity of these scattered lymph nodes did not change on qualitative readout, which was confirmed by quantification.

### Capture of circulating ^64^Cu-labeled OMVs by the spleen and liver after an intravenous administration

After an intravenous administration, the phagocytic system of the spleen and liver took up almost all ^64^Cu-labeled OMVs; however, the impurities of [^64^Cu]Cu-NOTA-N_3_ were immediately excreted via the kidneys and bladder, and the liver appeared to degrade the ^64^Cu-labeled OMVs (Fig. 2A, 2B; Suppl. Movie 4). The degradation products of ^64^Cu-labeled OMVs were traced with sequential PET imaging, and we identified two methods of excretion. The first method involves the use of [^64^Cu]Cu-NOTA or [^64^Cu]Cu-NOTA peptides, which are soluble and excreted via the kidneys and bladder; this process continues until the next day.

With this excretion of soluble ^64^Cu-labeled metabolites, another group of metabolites associated with cholic acids were excreted into the intestines (1-to 8-hour images) and the rectum (8-hour image). After 24 hours, radioactivity was continuously maintained by the phagocytic system of the spleen and the liver.

### Biodistribution of ^64^Cu-labeled OMVs after an intrathecal administration

We focused on three points regarding intrathecal administration. First, we focused on the volume, rate, and route of administration. In accordance with our previous reports (10, 16), we administered the sample in a 6-μL volume at a rate of 700 nL/min via the L5/6 disc space. Second, to adjust the administration volume and desired radioactivity, we pre-labeled NOTA-N_3_ with ^64^Cu and labeled DBCO-OMVs using click reaction and did not perform a separation procedure to remove the unlabeled [^64^Cu]Cu-NOTA-N_3_. The equivalents were optimized to yield 95% or higher labeling efficiency via instant liquid radiochromatography. Third, we investigated possible drainage, circulation, recirculation, metabolism and excretion or no absorption of ^64^Cu-labeled OMVs and their impurities. Notably, [^64^Cu]Cu-NOTA-N_3_ and rare hepatic degradation are important. We used intrathecal administration. This mimicked oral administration, as we administered the radiotracer into the seemingly closed compartment, very slow drainage from the CSF space and un-perturbing administration of the radiotracer made the situation physiologic and apparently in a closed compartment.

Immediately after the intrathecal administration, radioactivity was detected around the spinal cord; at 1 hour, it reached the basal cistern and dorsal parts of the cistern; at the paravertebral lymph nodes from the bilateral lower (deep) cervical, gastric/pancreaticoduodenal, lumbar aortic, medial iliac, and bilateral sacral levels (Fig. 3A, 3B; Suppl. Movie 5). The radioactivity of the eight lymph nodes and bilateral superficial cervical (also known as mandibular) lymph nodes, i.e., ten lymph nodes in total, appeared in the 1-hour images and was sustained throughout the 2-to 24-hour images (Fig. 3C). In terms of lymph node uptake, this mouse was representative; the other mice presented a similar pattern with a slightly delayed appearance of the upper abdominal lymph nodes, a delayed peak of the cervical lymph nodes (larger radioactivity in the 2-hour image than in the 1-hour image) and no contralateral sacral lymph node. The consistency of deep and superficial cervical uptake, both bilateral and bright, was prominent.

**Figure 3.**
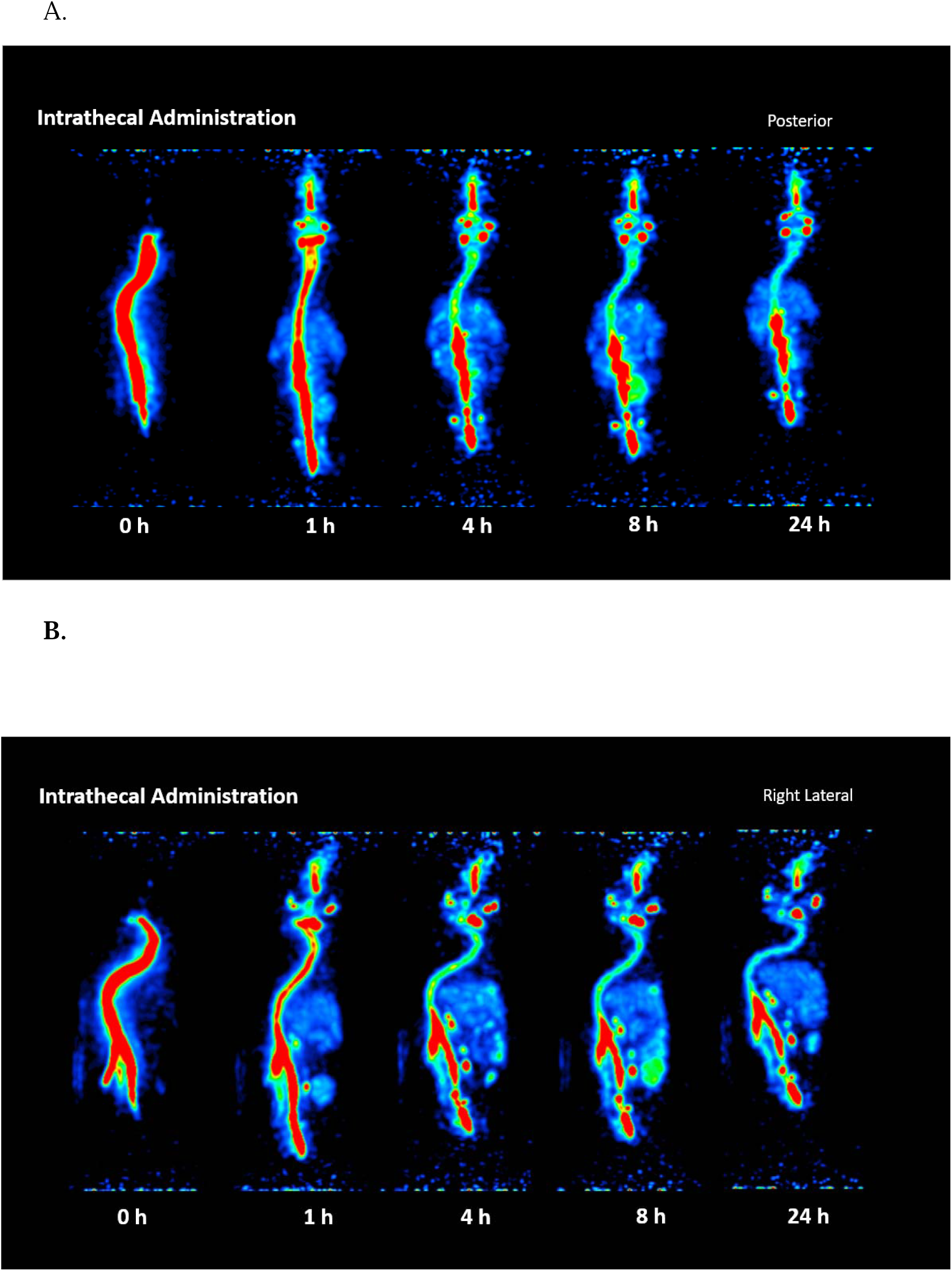
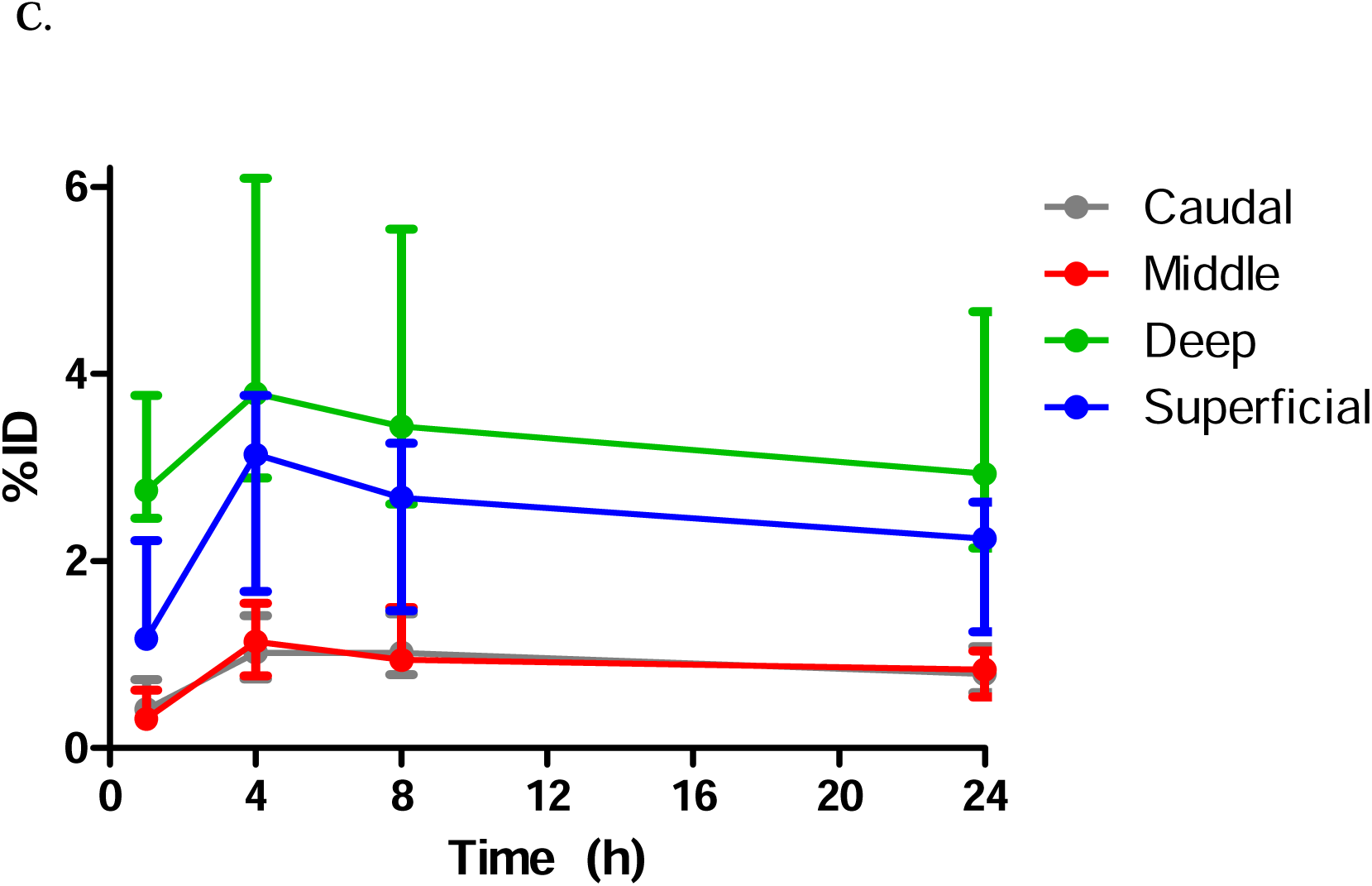
Biodistribution of ^64^Cu-labeled OMVs after the intrathecal administration in a mouse: A) posterior and B) right lateral views and C) quantification results in each segment of the lymph nodes. The bilateral deep cervical and sacral lymph nodes and the gastric/pancreaticoduodenal and lower lumbar/medial iliac nodes were considered paravertebral lymph nodes. The superficial lymph nodes were also bilaterally observed (C57BL/6 male, 7 weeks old). In three mice without leakage, as presented in Supplementary Table 1, the decay-corrected radioactivity of the segmental lymph nodes was drawn for the caudal (lower lumbar, medial iliac, and both sacral), middle (pancreaticoduodenal and gastric), deep (both deep cervical) and superficial (both superficial cervical) lymph nodes.

The bodily systemic faint uptake showed a similar pattern. Initially, there was no radioactivity in the bladder, which implies that the impurity [^64^Cu]Cu-NOTA did not appear in the bladder at 1 hour in all mice. In all three mice, radioactivity slightly appeared in the liver and subsequently in the intestine. It also faintly but continuously appeared in the bladder. This finding was considered to show the drainage of ^64^Cu-labeled OMVs after remaining/saturating in paravertebral/superficial cervical lymph nodes. ^64^Cu-labeled OMVs drained via the lymphatic channels, entered the systemic circulation and reached the liver to be slowly metabolized and excreted via the kidneys and hepatobiliary tract. In a mouse, the radioactivity was quite prominent in the liver and intestine, including the rectum, at 8 hours, which was reminiscent of the oral administration case.

The quantified results revealed a segmental distribution of drainage: upper (superficial and deep cervical 29.3% and 48.0%, respectively), 77.3%; middle (pancreaticoduodenal, gastric, lumbar aortic, and medial iliac), 11.4%; caudal (sacral), 11.3%. The appearance time or buildup and sustained radioactivity imply that all initial lymph nodes were the first stop of the drained ^64^Cu-labeled OMVs. Considering the behavior of the lymphatic-lymph node flow after a subcutaneous/intramuscular administration, the capture/keeping mechanism of ^64^Cu-labeled OMVs enabled the visualization of the first stop from the CSF of the brain and spinal cord.

### Biodistribution of ^64^Cu-labeled ENVs after intravenous and intrathecal administrations

We questioned whether the drainage and capture phenomenon was a common characteristic of EVs regardless of whether they originate from prokaryotes or mammalian cells. Whether the OMVs and mammalian EVs, especially those captured by the lymph nodes, have different biodistributions.

To answer this question, we produced ENVs as EV surrogates as described in (23) via a scalable high-shear microfluidic system from the isolated nucleus-free cellular membranes of immortalized human bone-marrow-derived mesenchymal stem cells (Fig. 4A). The produced ENVs were approximately 85.1 (±21.6) nm in diameter on average (Fig. 4B). SDS-PAGE showed that ENVs had different protein compositions from their parental cells (Fig. 4C). In the immunoblotting analysis, CD81, which is a well-known mammalian EV surface marker, was highly enriched in the purified ENVs compared to the whole-cell lysate. In addition, beta-actin (a cytosolic protein) and histone H2B (a nuclear protein) were substantially depleted in the purified ENVs compared to whole-cell lysates (Fig. 4D). The purified ENVs were labeled with ^64^Cu using the same method as the OMVs, as shown in Fig. 1A, and they were labeled with an efficiency equivalent to that of the OMVs.^64^Cu-labeled ENVs were intravenously administered to mice; then, the distribution pattern was determined in comparison with that of ^64^Cu-labeled OMVs (Fig. 5A, 5B; Suppl. Movie 6). After the intravenous administration, ^64^Cu-labeled ENVs circulated and left the impure [^64^Cu]Cu-NOTA to the kidneys and bladder. At 1 hour, ^64^Cu-labeled ENVs were retained in the liver and were excreted via the hepatobiliary tract. The radioactivity appeared in the gall bladder and was subsequently excreted when the mouse was fed *ad libitum*. The radioactivity appeared at 1 hour and 4 hours, faded and finally moved to the rectum at 8 hours. The radioactivity decreased in the liver at 24 and 48 hours. Note that the radioactivity never appeared in the spleen at various time points after the intravenous administration.

**Figure 4.**
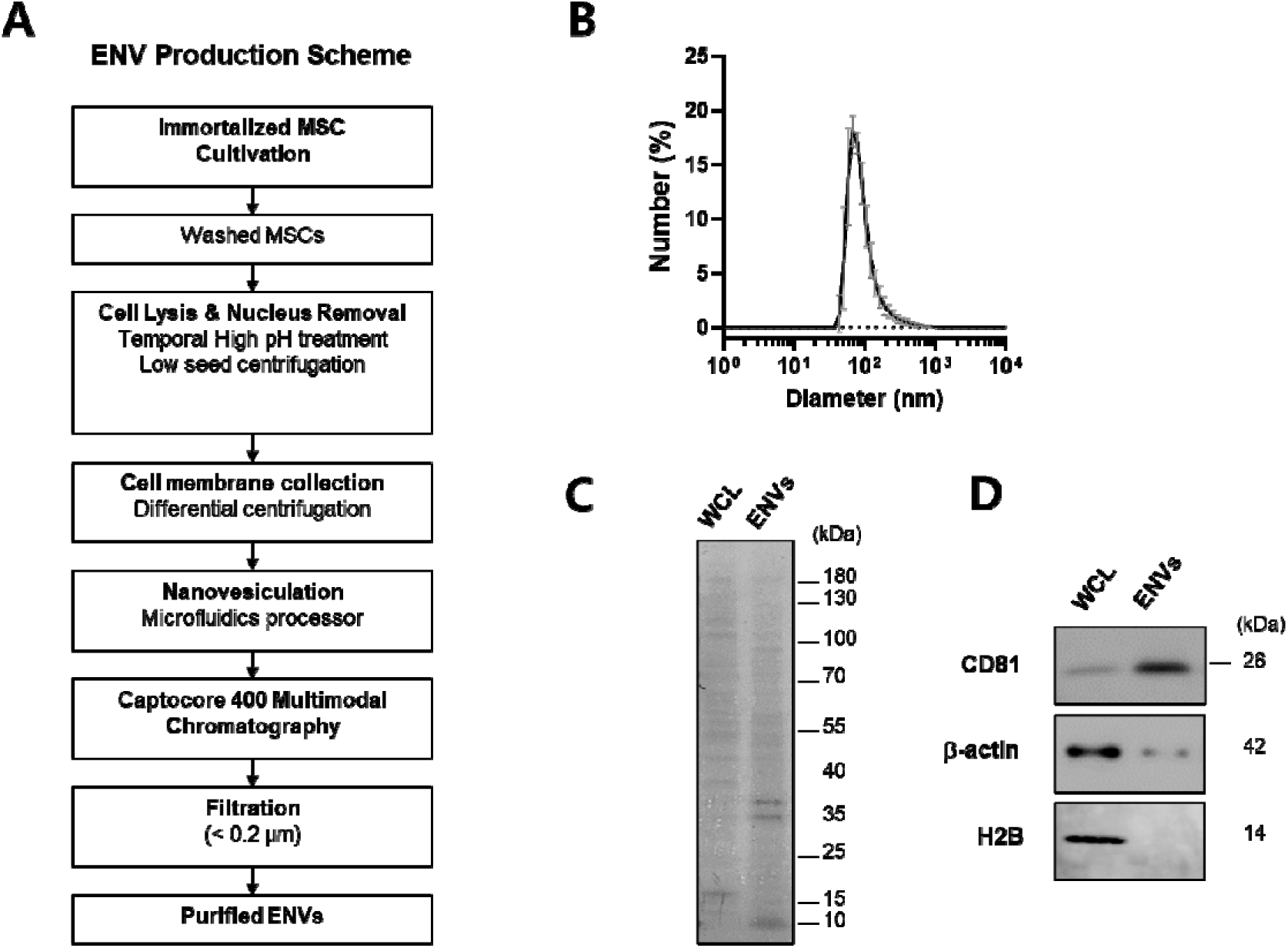
Production and characterization of ENVs. **(A)** Schematic of the ENV production. **(B)** Size distribution of ENVs measured using dynamic light scattering. The average size of the ENVs was 85.1 (±21.6) nm in diameter. **(C)** Total protein staining of ENVs and whole-cell lysates (protein amounts: 2 µg/well). **(D)** Immunoblot analysis of ENVs against CD81, ß-actin, and histone H2B protein (protein amount: 1 µg/well).

**Figure 5.**
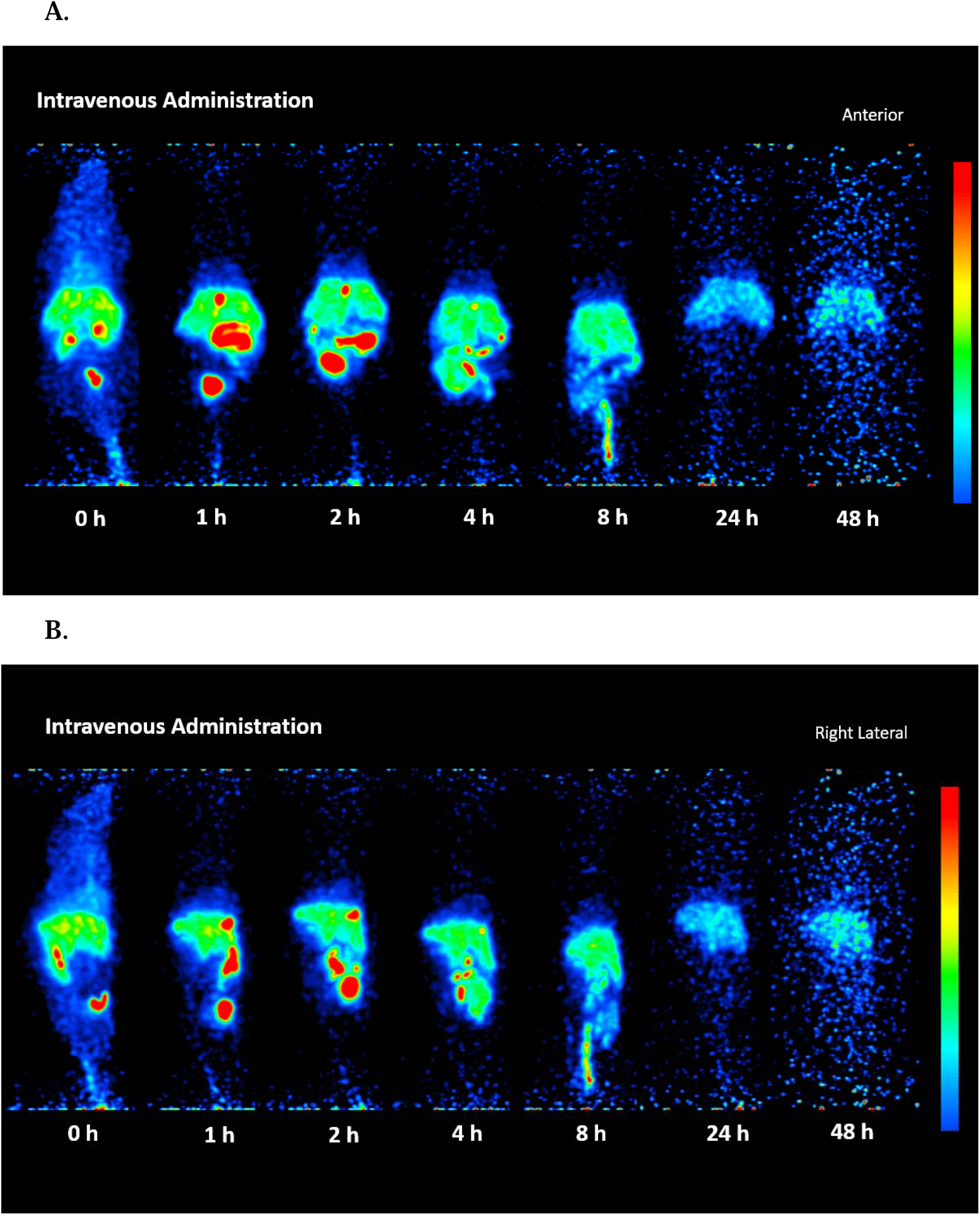
Biodistribution of ^64^Cu-labeled ENVs after intravenous administration in a mouse: A) anterior and B) right lateral views. ^64^Cu-labeled ENVs in the liver, and hepatobiliary excretion from the gallbladder, intestines and rectum. Initially, the kidneys excreted unlabeled [^64^Cu]Cu-NOTA-N_3_ up to 2 hours after the administration (BALB/c, male, 18 weeks old).

After the intrathecal administration, ^64^Cu-labeled ENVs were distributed throughout the entire spinal cord, with a small but definitely detectable distribution in the bladder (Figs. 6A, 6B; Suppl. Movie 7). At 1 hour, they reached the basal cistern, space around the olfactory bulb, superficial cervical lymph nodes, gall bladder, intestine and urinary bladder.

**Figure 6.**
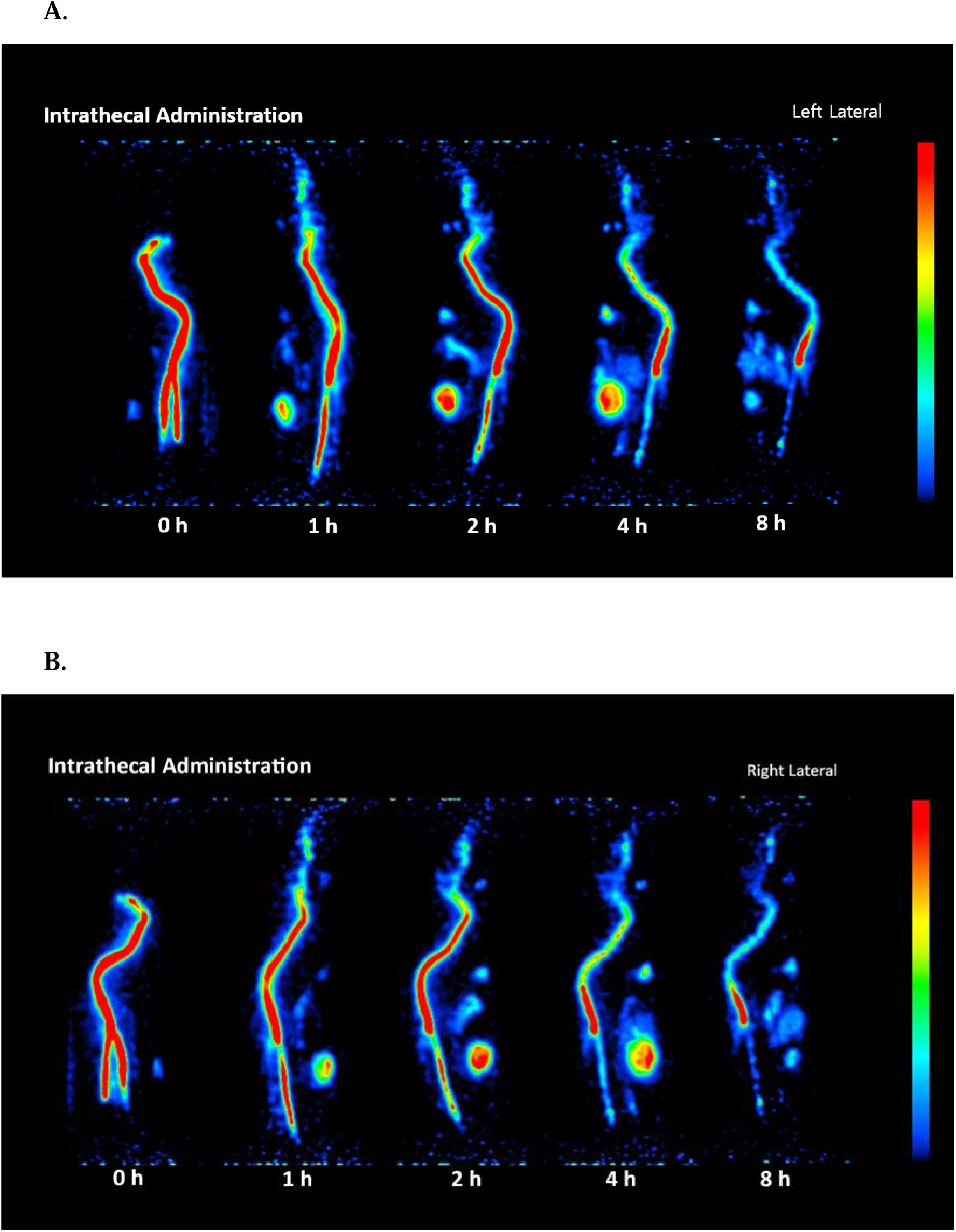
Biodistribution of ^64^Cu-labeled ENVs after the intrathecal administration in a mouse: A) left lateral and B) right lateral views. No paravertebral lymph nodes were observed in any image, and only superficial lymph nodes were observed. The bladder, gallbladder and intestinal activities revealed that the ^64^Cu-labeled ENVs in the CSF drained into the lymphatic system and circulated through the blood vascular system. Hepatic metabolism yielded [^64^Cu]Cu-NOTA fragments, which were partly excreted via the kidneys and hepatobiliary tract (BALB/c male, 18 weeks old).

Interestingly, radioactivity of ^64^Cu-labeled ENVs did not appear in the liver. It did not either in paravetebral lymph nodes despite the similar remaining amounts of ^64^Cu-labeled OMVs at 8-hour image. These labeled ENVs were drained to the systemic circulation, considering the total bodily activity, however in different pattern of passing lymphatics and lymph nodes from that of ^64^Cu-labeled OMVs. ^64^Cu-labeled ENVs in CSF would have reached the liver, which metabolized ^64^Cu-labeled ENVs to [^64^Cu]Cu-NOTA/[^64^Cu]Cu-NOTA-peptides and [^64^Cu]Cu-NOTA-larger fragments with the lipid components. Among these metabolites, [^64^Cu]Cu-NOTA/[^64^Cu]Cu-NOTA-peptides were rapidly cleared via the kidneys to fill the bladder and removed as urine. [^64^Cu]Cu-NOTA-larger fragments with lipid components are excreted via bile acid assistance to the intestines and finally to the rectum/outside the body. Superficial cervical lymph nodes are the only evidence that ^64^Cu-labeled OMVs passed through the CSF-lymphatic drainage pathway. Nevertheless, ^64^Cu-labeled ENVs in CSF appeared to drain via the same route as ^64^Cu-labeled OMVs, considering that the radioactivity in CSF upon intrathecal administration decreased at a similar rate, the radioactivity appeared in the body other than the brain and spinal cord.

## Discussion

In this *in vivo* imaging experiment, we investigated the location and function of paravertebral lymph nodes to monitor the brain and spinal cord in mice. Ten lymph nodes, to which ^64^Cu-labeled OMVs and ENVs were intrathecally administered around the lower lumbar CSF space without spillage, were the first stop. This finding was obtained using two characteristics of PET imaging; 1) ^64^Cu-labeled OMVs could be caught by the lymph nodes that met for the first time within the lymphatic vessels; 2) ^64^Cu-labeled OMVs or ENVs administered in a compartment will release radioactive emission, which can be imaged with ultimate contrast (infinite contrast) in cases of oral, subcutaneous, intramuscular, and most importantly intrathecal routes but not intravenous route. Interestingly, the intravenous administration of ^64^Cu-labeled OMVs and the use of ^64^Cu-labeled ENVs played the roles of supplying lessons about the metabolism/excretion and counterpart controls as a negative substance for visualization, respectively. We currently believe that the CSF is drained within this amount (fraction) to the lymph nodes of the cervical level, lower thoraco-abdominal level, and lumbo-sacral level. Individual variation could be documented in separate mice. Thus, this method is for individual animal study.

The first advantages of radiolabeled EVs were first reported by us using luminal [^99m^Tc]Tc-HMPAO labeling of ENVs (24,25). Then, the following trials focused on the lipid membrane labeling including our own failing to follow the long-term fates of EVs or ENVs *in vivo* (24–28). Non-covalent bond labeling has been known to cause problem in tracing the biodistribution of EVs or ENVs in its integrity. Among them, fluorescent dye was the best known as the detached dye permanently stays in the body and show its own distribution (26–28). Brain visualization was the most cumbersome for this. Iodine or other mimetics ([^99m^Tc]TcO^4-^) was also for thyroid visualization (29–31). Luminal [^99m^Tc]Tc-HMPAO labeling of EVs or ENVs also posed challenges for image interpretation. Since the luminal metabolite of [^99m^Tc]Tc-HMPAO is hydrophilic, once the EV/ENV is leaked, it follows the hepatobiliary excretion pathway through the liver and intestines (24). This hepatobiliary excretion pathway of hydrophilic metabolite of [^99m^Tc]Tc-HMPAO was identical to the metabolized ^64^Cu-labeled ENVs and ^64^Cu-labeled OMVs of this investigation. The difference was that the metabolized products of the OMV/ENV membranes tightly held the [^64^Cu]Cu-NOTA-peptide fragments and selected their route to excrete via hepatobiliary or renal pathways. We adopted the click reaction of alkyne-azide and prelabeled [^64^Cu]Cu-NOTA-N_3_ (20). The reaction was almost complete with a small amount of unlabeled [^64^Cu]Cu-NOTA-N_3_, which made the after-labeling separation procedure unnecessary. On the other hand, the small amount of unlabeled [^64^Cu]Cu-NOTA-N_3_ yielded the chances to understand and interpret the sequential biodistribution of originally unlabeled [^64^Cu]Cu-NOTA-N_3_ and the metabolized [^64^Cu]Cu-NOTA-N_3_ with or without peptide fragments and their differences.

OMVs were developed as cancer immunotherapeutic agents by Gho’s group (17) and later mass-produced OMVs in the following report to be used as stimulant for CD8+ T cell in combination cancer immunotherapy (18). Intra-tumoral routes were favored as booster and the whereabouts of OMVs raised the curiosity. We examined this question in the present investigation and found that most of the subcutaneously administrated ^64^Cu-labeled OMVs stayed within injection sites and small amount revealed the lymphatic outflow from muscles of subcutaneous spaces (Suppl. Fig. 2AB, Suppl. Movies 2AB). We observed that the clearance patterns of ^64^Cu-labeled OMVs varied depending on the route of administration, likely due to the differences in lymphatic drainage pathways for each route. Subcutaneous injections enabled broader diffusion through the superficial lymphatic system and caused drainage into nearby and distant lymph nodes, whereas intramuscular injections showed more localized absorption with faster entry into the bloodstream due to the rich vascularization of muscle tissue. More diffuse but significant drainage to the blood vessels via thoracic and right lymphatic duct (32) were followed by the hepato-biliary (gall bladder) excretion to the intestines. The drained lymphatics of the intramuscular site carried ^64^Cu-labeled OMVs to the first (sentinel) lymph node and secondary ones. This lymph-node uptake characteristics was expected and proven in this comparison study of the fates of ^64^Cu-labeled OMVs using various routes. The imaging finding of one route reciprocally and directedly supported another. The lymph node uptake characteristics of ^64^Cu-labeled OMVs were finally contrasted with the absence of this characteristics of a mammalian (human) mesenchymal stem cell-derived ^64^Cu-labeled ENVs; this is another example that every group of EVs are different in their behavior *in vivo* (33–35). Thus, we speculate that ^64^Cu-labeled OMVs could not be used for drug delivery via oral route. This speculation contradicts the report on the systemic uptake of orally administrated DiD-labeled OMVs (36), which might have been due to fluorescent dye released from OMVs (26,27,37). We suspect that the intestinal luminal space is full of OMVs and EVs derived from various microbiome and host itself and added trace amount of ^64^Cu-labeled OMVs were released via esophago-gastro-intestinal-colic-rectal peristalsis (Suppl. Fig. 1, Suppl. Movie 1). Two items must be clarified: gastric radioactivity stayed too long, though evacuated at last 2 days after, considering minutes of gastric liquid emptying and tens of minutes of gastric solid emptying.

Intrathecal administration was prominent in taking advantage of nuclear imaging*, i.e*., click reaction-labeled ENVs/OMVs PET, compartmental injection and follow-up for tracing the fate at the lymph node stopping OMVs *in vivo*. Here, we visualized and quantified the CSF-lymphatic drainage in mice. The clearance of CSF radioactivity of OMVs was slower than albumin similarly labeled with click reaction (Suppl. Fig. 4) (10). The lymph nodes as the first stops were deep cervical lymph nodes (bilateral), pancreaticoduodenal and gastric (one for each), lower lumbar aortic (one at the midline), medial inguinal (one at midline) and sacral (bilateral) lymph nodes (38–40). There were eight in total. In addition, superficial cervical lymph nodes (bilateral) participated. Individual variation could be explained by the amount of total administered volume of intrathecal ^64^Cu-labeled OMVs and their fraction of the CSF of each mouse. The rate was exactly controlled, and the total amount was identical (10,16), but when we scrutinized the early images, the time of arrival of radioactivity to the basal cistern and the paravertebral lymph nodes differed mouse to mouse. Another individual variation was the unilateral non-visualization of the sacral lymph node and relatively varying intensity of the abdominal lymph nodes. In contrast, superficial and deep cervical lymph nodes showed consistent symmetric radioactivity and ratio of radioactivity of both lymph nodes over the radioactivity of CSF space.

We present our findings in the context of the well-established connections among the brain, spinal cord, and lymphatic and circulatory systems of the body. The skull and vertebrae have bone marrow, whose marrow sinus veins are connected with the dura sinus veins via bridging veins (41). Dura veins are connected with the brain parenchyma via other bridging veins (13). The arachnoid barrier cell layer is tightly closed with claudin-11; an exception is the structured gaps, which used to be called cranial arachnoid granulation-like dural gaps (CAG-DGs) in pigs (11) and subsequently found in mice as defects in the arachnoid barrier cell sheets between dura vein and subarachnoid space (12). This gap/defect was again reported in both mice and humans and is called the arachnoid cuff exit (13). We now know that the subarachnoid space CSF is not entirely separated from the dura extracellular matrix, and using these gaps/defects/exits of fluid, small molecules, macromolecules, and particulate materials such as EVs can move out of the brain and spinal cord. Routinely and/or if needed, skull/vertebrae bone marrow cells (mostly immune) can commute to and from the brain/spinal cord parenchyma and subarachnoid CSF spaces (42,43).

Fluorescent dyes labeled with various sized materials (4,5), radio-labeled inulin/proteins (16,44,45), and MRI contrast dyes (46,47) were successfully used to visualize or quantify the drainage of CSF, probably via the lymphatic system. The investigators dedicated a lot of effort to understand the condition and obstacles or perturbation effects upon these normal CSF-lymphatic drainage systems. Here, we used a lymph node seeker, ^64^Cu-labeled OMVs (Suppl. Fig. 2 and 3; Suppl. Movies 2AB and 3), and an established radiochemistry and animal PET system to identify the first stop of CSF-lymphatic drainage in mice. The optimal amount/rate of ^64^Cu-labeled OMVs was intrathecally administered, circulated throughout the cisterns and peri-spinal cord CSF spaces, and casually drained to the regional lymphatics via adjacent lymphatics (10,16). ^64^Cu-labeled OMVs in lymphatic vessels that emanate from inside the arachnoid barrier cell layer around the brain and spinal cord are destined to be captured by the lymph nodes; they remain there and indicate that they originated from CSF.

Based on these observations concerning the situation of the lymph nodes and central nervous system, we propose that the brain and spinal cord release waste via the CSF to the lymphatic system, which benefits the gaps/exits/defects of arachnoid barrier cells. Draining lymphatics, which were visualized using fluorescent dye-labeled ovalbumin injected into the intrathecal spaces and subsequently found in the epidural lymphatics, gathered together with regional lymph nodes via valves and/or lymphatic vascular contraction and hydrostatic pressure differences. Due to an unknown mechanism, the lymph nodes only caught the OMVs but not the other substances. Drained lymphatic was recently clarified to move via nasopharyngeal lymphatic plexus and easily reach the superficial or deep cervical lymph nodes (15). The perineural sheaths of cranial nerves have been proposed as routes for CSF to move through systemic lymphatic channels outside the brain/spinal cord (48,49). However, the cribriform plates with olfactory nerves have been reported to be the main routes with many submucosal plexus forms in the vast nasopharyngeal area (15), whereas the dorsal dura meningeal lymphatics also drain CSF waste.

We suggest that the CSF-lymphatic drainage follows the general rule of interstitial space-lymphatics interaction, i.e., “demand is accompanied by drainage”. The demand follows the supply and surface area of the dura of the brain and spinal cord. The waste supplied from the brain and spinal cord demands removal from the CSF, which is necessary for the surface to pass through. Despite the absence of the measured dural surface area, we estimated the surface area of the brain and spinal cord using their reported weights (50,51). The weights of the male C57BL/6 brain and spinal cord were 488 mg and 132 mg, respectively, whose surface areas were converted to 62 mm^3^ and 26 mm^3^, respectively. The surface area of the brain was 70% of the total brain and spinal cord area, which was similar to the 77% of uptake among the administered ^64^Cu-labeled OMVs at the cervical lymph nodes. We propose that the cervical lymph nodes receive drainage from the cervical spinal cord in addition to the brain. In addition to the demand, the local condition of the lymph nodes themselves, which is related to the position, and probable latent inflammation may have affected the drainage proportion. The abdominal and pelvic lymph nodes varied the most (16,48, Suppl. Table 1), and the sacral lymph nodes showed definitive asymmetry (Fig. 3A, Suppl. Table 1, Suppl. Movies 7,8,9). This large variation, including sometimes non-visualization of one side, is the reality of the CSF-lymphatic drainage of each individual mouse. Individual variation should be expected in any mouse; otherwise, normal states of CSF-lymphatic drainage are presumed to vary.

The mechanism by which OMVs affect the lymph nodes remains to be determined. The variation in the lymph node-seeking ability of other bacterial or prokaryotic OMVs or EVs is also unknown. Mammalian or other human cell-derived EVs or ENVs of various sizes are interesting. We are certain that the highly purified and preserved EVs, ENVs, and OMVs should be labeled with chelator pre-bound ^64^Cu or other relevant radionuclides. No separation procedures are preferred to preserve the specificity of labeling (Bq/mol) and not to damage the EVs, ENVs, or OMVs. Serial imaging is recommended, or at least delayed imaging is mandatory. Since membrane binding is very tight and robust in every route of administration, we recommend that investigators interpret the linear rectum radioactivity in oral administration, sub-skin faint but diffuse radioactivity in subcutaneous administration, and gallbladder radioactivity in the hepatobiliary excretion and differential initial renal/bladder radioactivity or delayed/sustaining bladder activity after the hepatic metabolism. The most important factor was the degree of lymph node radioactivity. We suggest that all lymph node activities indicate the first stop, especially from the CSF space, after the intrathecal administration. This imaging method can now be used to evaluate the effect of any disease-modifying treatment in mice to determine whether these treatments improve the CSF-lymphatic drainage. If these works are reproduced, the CSF-lymphatic drainage can also be easily evaluated after drainage modulation and/or stimulation (52–54).

## Methods

### Mice

Seven-week- or 14-week-old C57BL/6 and 18-week-old BALB/c male mice were used for imaging. All experimental protocols were approved by the Institutional Animal Care and Use Committee (IACUC) at Seoul National University (SNU211019-1 and SNU-230802-3). All methods were performed according to relevant IACUC guidelines and regulations of Seoul National University and ARRIVE guidelines (https://arriveguidelines.org).

### Preparation of OMVs and ENVs

#### *E. coli* OMV purification

We used purified OMVs as previously reported (18). Briefly, the *E. coli* BL21(DE3) ΔmsbB mutant (SL Bigen Co. Ltd., Korea) was cultured in particle-free vegetable lysogeny broth at 30°C for 16 hours. The cells were removed via centrifugation at 10,000 ×g, and the cell-free culture medium was subsequently filtered through a 0.2-μm filter. The clarified conditioned medium was concentrated using a tangential flow filtration system with a 100-kDa molecular weight cutoff (MWCO). OMVs in the concentrated medium were enriched via metal-based precipitation. The resulting crude OMVs were subjected to size exclusion chromatography using Sephacryl S300 (Cytiva, USA). Then, the purified OMVs were filtered through a 0.1-μm filter and stored at -80°C until use.

#### ENV Production

BM300, which is an immortalized human bone marrow-derived MSC that expresses CD47 on its surface (SL Bigen, Co. Ltd., Korea), was cultured in Dulbecco’s Modified Eagle Medium (Gibco, USA) supplemented with 10% fetal bovine serum (FBS) and 10 ng/mL recombinant human fibroblast growth factor-basic under humidified 5% CO_2_ at 37°C. The harvested cells were ruptured in a solution that contained 1 mM Na[CO[ and 1 mM MgCl[. The nuclear compartments were removed by centrifugation at 1,000 *× g*. Then, the membrane fraction was collected by high-speed centrifugation at 12,000 *× g* for 30 minutes. To generate ENVs, the isolated cellular membranes were subjected to shear-induced nanovesiculation via an LM10 high-shear microfluidic process (Microfluidics^TM^, USA) at 18,000 psi. The resulting ENVs were further purified via multimodal column chromatography using Captocore^TM^ 400 (Cytiva, USA). Then, the purified ENVs were filtered through a 0.2-µm filter and stored at -80°C until use.

### Characterization of ENVs

#### Dynamic light scattering analysis

The size distribution of ENVs was measured using a Zetasizer Nano ZS (Malvern Instruments Ltd. UK). The ENVs (10 μg/mL each) were subjected to the following parameters: laser wavelength = 633 nm; number of measurements = 5; measurement duration = 30 sec; chamber temperature = 25°C.

#### SDS-PAGE and immunoblotting analysis

The equivalent protein amounts of whole-cell lysates and ENVs were denatured and loaded into each well of 4-20% polyacrylamide gels (Invitrogen, USA). After the electrophoresis, the gels were subjected to SimplyBlue protein staining (Invitrogen, USA) to visualize the proteins. For the immunoblotting analysis, the gels were transferred to a polyvinylidene difluoride membrane. The blots were blocked with 5% skim milk and incubated with mouse monoclonal anti-CD81 (555875, BD, USA), anti-β-actin (sc-47778, Santa Cruz Biotechnology, USA), or rabbit polyclonal anti-H2B antibody (07-371, Millipore, USA). Subsequently, the blots were incubated with horseradish peroxidase-conjugated secondary antibodies, and the immune complexes were visualized via incubation with an enhanced chemiluminescence substrate (Thermo Fisher Scientific, USA).

### Click chemistry labeling of ^64^Cu to OMVs and ENVs and the radio-labeling efficiency

OMVs and ENVs were radio-labeled with ^64^Cu using an azide-alkyne click chemistry-based method. Briefly, the ^64^Cu radionuclide-NOTA-N_3_ complex was prepared by a selective coordination of ^64^Cu via a NOTA-N_3_ (3-azidopropyl-NOTA) chelating agent.

OMVs or ENVs were incubated with 3 molar excesses of DBCO-NH-ester at pH 8.3 for 1 hour at room temperature. After 1 M ethanolamine had been added to block the free ester, the unbound DBCO was removed via diafiltration using Amicon™ Ultra centrifugal filter devices with a 100-kDa MWCO (Sigma-Aldrich, USA). OMVs or ENVs were conjugated with DBCO-NHS ester in phosphate-buffered saline (PBS, pH 7.4) for 30 minutes at 25[. To remove the unlabeled DBCO-NHS ester from the OMVs or ENVs, the mixture was purified four times using centrifugal filter with a centrifuge at 4,550 _D_ g for 5 minutes at 25.

The HCl solution of ^64^Cu was heated at 70[ until every solution that contained ^64^Cu was evaporated by N_2_ gas for 30 minutes. After the ^64^Cu solution had been evaporated, the pH level was adjusted to 5 by adding 1 M sodium acetate buffer (pH 5). 3-Azidopropyl-NOTA was added to ^64^Cu, which was titrated to pH 5 and reacted at 80[ for 10 minutes. The radio-labeling efficiency was determined using radio instant thin-layer chromatography-silica gel (iTLC-SG) with 0.1 M citric acid as the mobile phase. [^64^Cu]Cu-NOTA-N_3_ was added to DBCO-labeled OMVs and ENVs, which were reacted at 4 for 16 hours. Radio-labeled OMVs and ENVs were analyzed using iTLC-SG with 0.1 M citric acid as the mobile phage to evaluate the radio-labeling efficiency.

### Imaging of ^64^Cu-labeled OMVs: Oral, subcutaneous, intramuscular, intraperitoneal and intravenous administration

Imaging of ^64^Cu-labeled OMVs was performed using GENISYS^4^ (SOFIE Biosciences). All animal studies were approved by the Institutional Animal Care and Use Committee at Seoul National University Hospital. Seven-week-old mice were scanned for 3 minutes via static scanning after subcutaneous, intramuscular, intraperitoneal and intravenous administrations of adequate ^64^Cu radioactivity that was suitable for administration routes (subcutaneous =2.50 MBq; intramuscular = 1.52 MBq; intraperitoneal =2.89 MBq; intravenous = 2.59 MBq). During the imaging studies, the mice were anesthetized with isoflurane containing 3% O_2_. All images were reconstructed using the internal algorithm of the GENISYS^4^ software.

### Imaging of ^64^Cu-labeled OMVs: Intrathecal administration

Imaging of ^64^Cu-labeled OMVs was performed using GENISYS^4^ (SOFIE Biosciences). This animal study was approved by the Institutional Animal Care and Use Committee at Seoul National University Hospital. Seven-week-old mice were scanned for 5 minutes via static scanning after an intrathecal administration of 0.97 MBq ^64^Cu. During the imaging study, the mice were anesthetized with isoflurane containing 3% O_2_. All images were reconstructed using the internal algorithm of the GENISYS^4^ software.

To quantify the segmental distribution of ^64^Cu-labeled OMV drainage from the CSF to various lymph nodes, we first measured the percentage of injected dose (%ID) that remained in the CSF at multiple time points (0, 1, 4, 8, and 24 hours) across three subjects. The decay of ^64^Cu in the CSF was modeled using an exponential function. Assuming that the ^64^Cu-labeled OMV input to each lymph node was proportional to the CSF output, we normalized the %ID uptake in the caudal, middle, and upper (deep and superficial) lymph nodes to the CSF output. Time[activity curves were generated for each segment of the lymph nodes, and the cumulative activity was used to calculate the percentage of total drainage from the CSF to each segment of the lymph nodes. The upper segment was subdivided into deep and superficial lymph node segments.

### Imaging of ^64^Cu-labeled ENVs: Intravenous and intrathecal administration

Imaging of ^64^Cu-labeled ENVs was performed using GENISYS^4^ (SOFIE Biosciences). All animal studies were approved by the Institutional Animal Care and Use Committee at Seoul National University Hospital. Seven-week-old mice were scanned during the appropriate time (intravenously = 3 minutes; intrathecally = 10 minutes) via static scanning after intravenous and intrathecal administrations of adequate ^64^Cu radioactivity, which were suitable for administration routes (intravenously = 2.66 MBq; intrathecally = 0.22 MBq). During the imaging studies, the mice were anesthetized with isoflurane containing 3% O_2_. All images were reconstructed using the internal algorithm of the GENISYS^4^ software.

## Author contributions

KK: Methodology; Investigation; Imaging; Writing – original draft.

CL: Methodology; Investigation; Imaging; Writing – original draft.

RJY: Methodology; Investigation; Imaging.

YC: Methodology; Investigation; Imaging.

CL: Methodology; Investigation.

JL: Writing – review & editing.

MS: Conceptualization; Methodology; Investigation; Imaging;

Writing – original draft; Writing – review & editing.

YSL: Conceptualization; Methodology; Investigation; Imaging; Writing – original draft;

Project administration; Supervision; Writing – review & editing.

YSG: Conceptualization; Methodology; Investigation; Imaging; Writing – original draft;

Project administration; Supervision; Writing – original draft; Writing – review & editing.

DSL: Conceptualization; Methodology; Investigation; Imaging; Funding acquisition; Project administration; Supervision; Writing – original draft; Writing – review & editing.

## Availability of data and materials

All data are available in the main text or supplementary materials. All data, code, and materials in the analysis are available through a standard material transfer agreement with POSTECH for academic and nonprofit purposes by contacting the corresponding authors.

## Supplementary Files

### List of Supplementary Figures

**Supplementary Figure 1. Biodistribution of ^64^Cu labeled OMVs after oral administration in mice.**

**Supplementary Figure 2. Biodistribution of ^64^Cu labeled OMVs after subcutaneous or intramuscular administration in mice.**

**Supplementary Figure 3. Biodistribution of ^64^Cu labeled OMVs after intraperitoneal administration in mice.**

**Supplementary Figure 4. Kinetics of intra-CSF space radioactivity %ID over the injected amount or ^64^Cu labeled OMVs after intrathecal administration in mice (n=3).**

**Supplementary Table 1. Ten lymph nodes and their uptakes of ^64^Cu labeled OMVs after intrathecal administration.**

**Supplementary Movie 1. Biodistribution of ^64^Cu labeled OMVs after oral administration in a mouse.**

**Supplementary Movie 2A. Biodistribution of ^64^Cu labeled OMVs after subcutaneous administration in mice.**

**Supplementary Movie 2B. Biodistribution of ^64^Cu labeled OMVs after intramuscular administration in a mouse.**

**Supplementary Movie 3. Biodistribution of ^64^Cu labeled OMVs after intraperitoneal administration in a mouse.**

**Supplementary Movie 4. Biodistribution of ^64^Cu labeled OMVs after intravenous administration in a mouse.**

**Supplementary Movie 5. Biodistribution of ^64^Cu labeled OMVs after intrathecal administration in a mouse**

**Supplementary Movie 6. Biodistribution of ^64^Cu labeled ENVs after intravenous administration in a mouse**

**Supplementary Movie 7. Biodistribution of ^64^Cu labeled ENVs after intrathecal administration in a mouse**

**Supplementary Movie 8. Biodistribution of ^64^Cu labeled OMVs after intrathecal administration in another mouse**

**Supplementary Movie 9. Biodistribution of ^64^Cu labeled OMVs after intrathecal administration in the third mouse**

**Supplementary Movie 10. Biodistribution of ^64^Cu labeled OMVs after intrathecal administration in the fourth mouse with leakage to epineural space.**

**Supplementary Figure 1.**
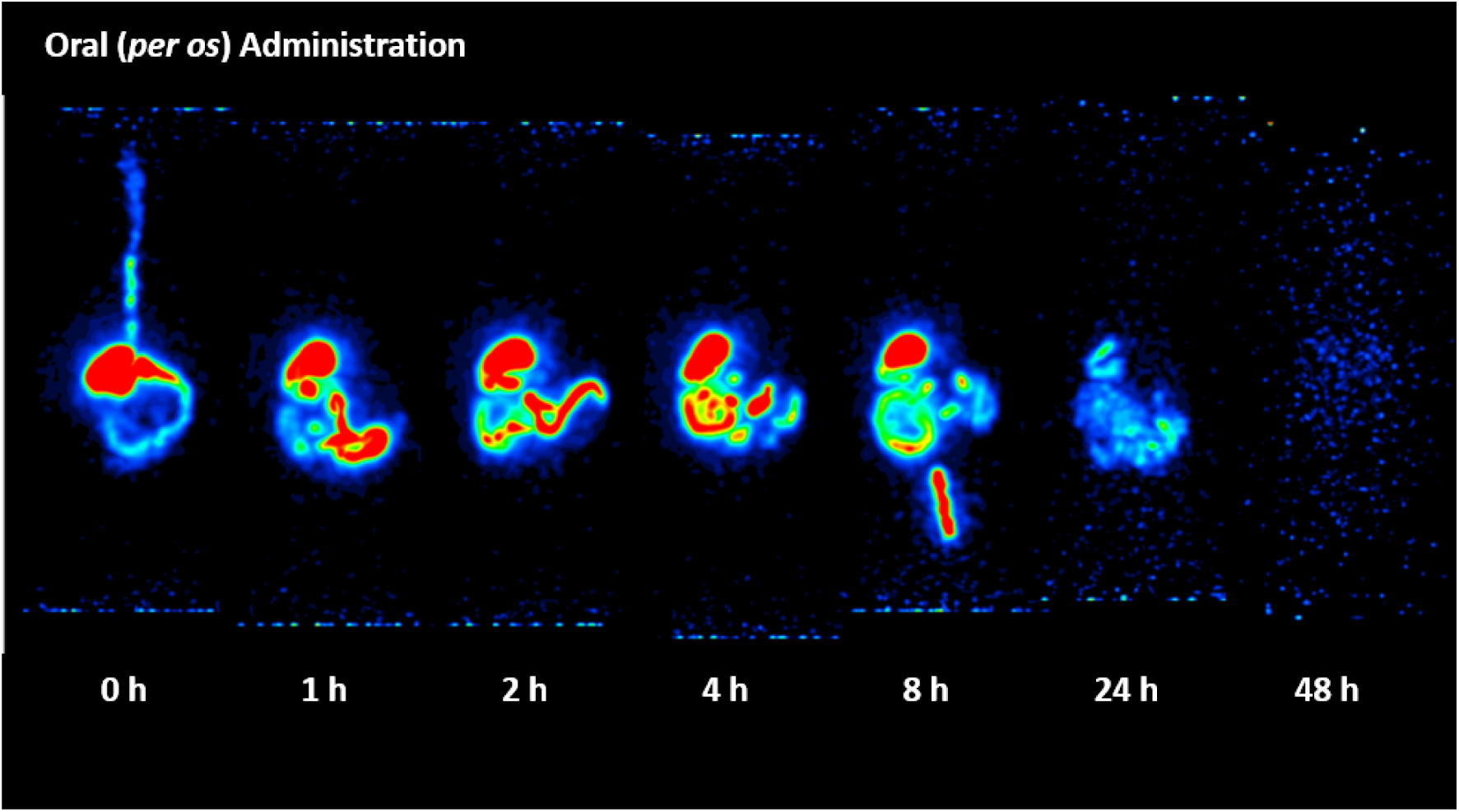
Biodistribution of ^64^Cu labeled OMVs after oral administration in mice. 48 hours after administration all the radioactivity is gone. 24-hour image showed the remnants which were scattered within stomach, small intestines and large intestines. We propose that the remaining OMVs were attached the gastrointestinal surface probably mucosa or microbiome and finally released to the lumen and outside of the body as feces. At 8 hour, rectum contained feces and the radioactivity. Stomach contained the radioactivity until 8 hours surpassing the expectation that the half time of liquid or solid food were minutes and tens of minutes. 8-hours were very long and the administered ^64^Cu labeled OMVs stayed a long time in the stomach beyond the transit time. The findings of this imaging is against the speculation that OMVs or their generic ENVs do not cross the mucosa-epithelial barrier of the gastrointestinal tracts. However, these images are not the evidence of the intra-OMVs or ENVs contents being taken up by the body, though the possibility is low.

**Supplementary Figure 2.**
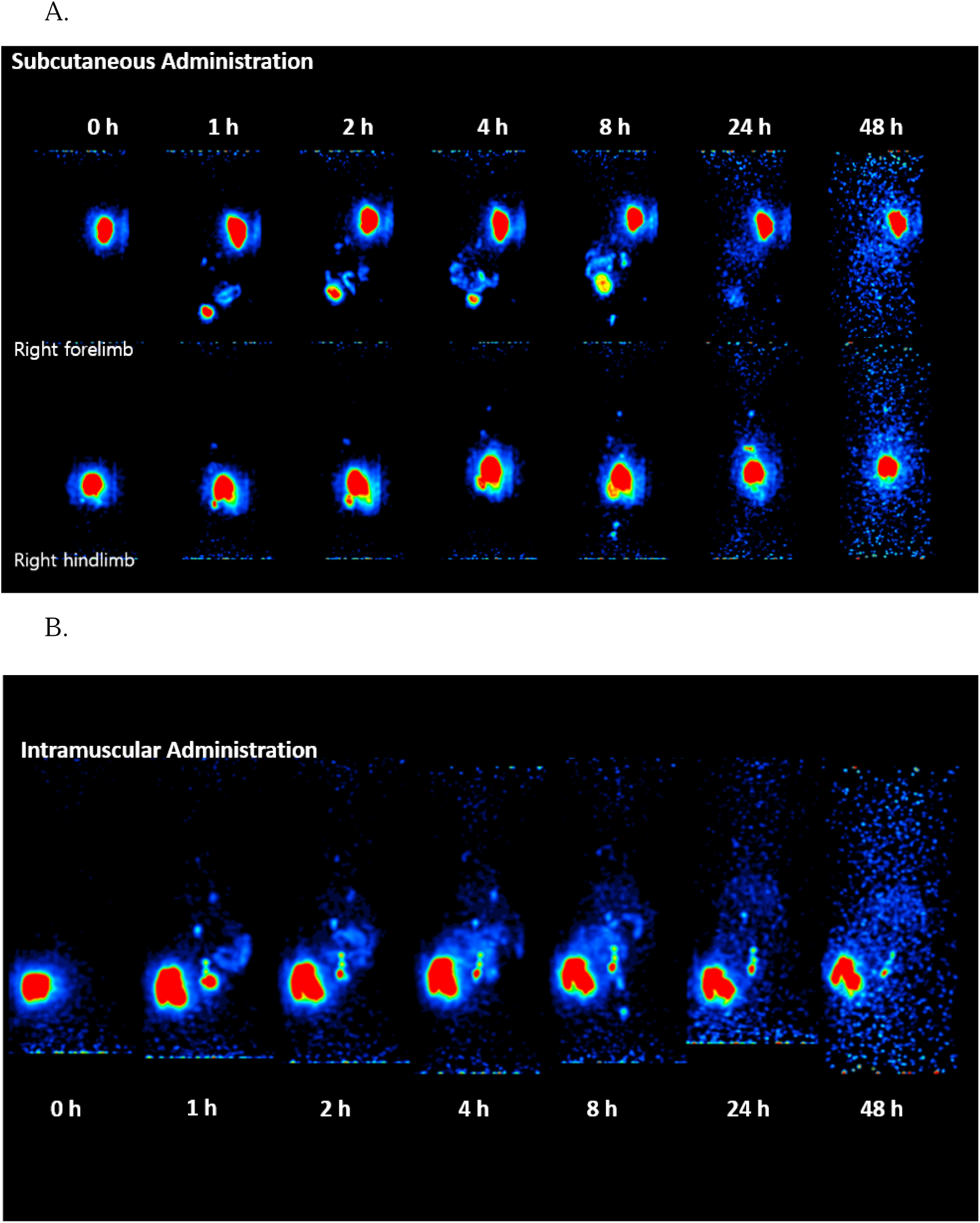
Biodistribution of ^64^Cu labeled OMVs after subcutaneous or intramuscular administration in mice. **A**) After subcutaneous injection of ^64^Cu labeled OMVs to the right hindlimb (upper row), radioactivity stayed on site for a long time till 48 hours. Faint distributed radioactivity appeared and persisted along the skin overlying the injection site suggesting dermal backflow. Bladder activity appeared at 1 hour in both hindlimb and forelimb (lower row) injections and continued till 8 hours, which disappeared the next day. Slow flow into the systemic blood circulation via the thoracic duct (mild but sustaining radioactivity at the inlet in hindlimb case) made intestinal activity, suggesting hepatic excretion, down to rectum (8-hour image). **B**) After intramuscular injection of ^64^Cu labeled OMVs to the right hindlimb, localized radioactivity stayed there on site until 48 hours as after subcutaneous injection. However, bladder radioactivity was different in that the radioactivity appeared at 1 hour imaging and then disappeared. We interpret this phenomenon that ^64^Cu labeled OMVs in the muscles stay on site but that impurity [^64^Cu]Cu-NOTA-N_3_ is quickly drained to lymphatics (deeper than subcutaneous layer) to reach thoracic duct and blood circulation to be evacuated via kidneys and bladder. After 1 hour no [^64^Cu]Cu-NOTA-N_3_ is inside the body. Interestingly, drainage of ^64^Cu labeled OMVs was more localized and prominent and sequential, which looked like a chain of lymph nodes, i.e., first lymph node brighter and then dimming to the third one.

**Supplementary Figure 3.**
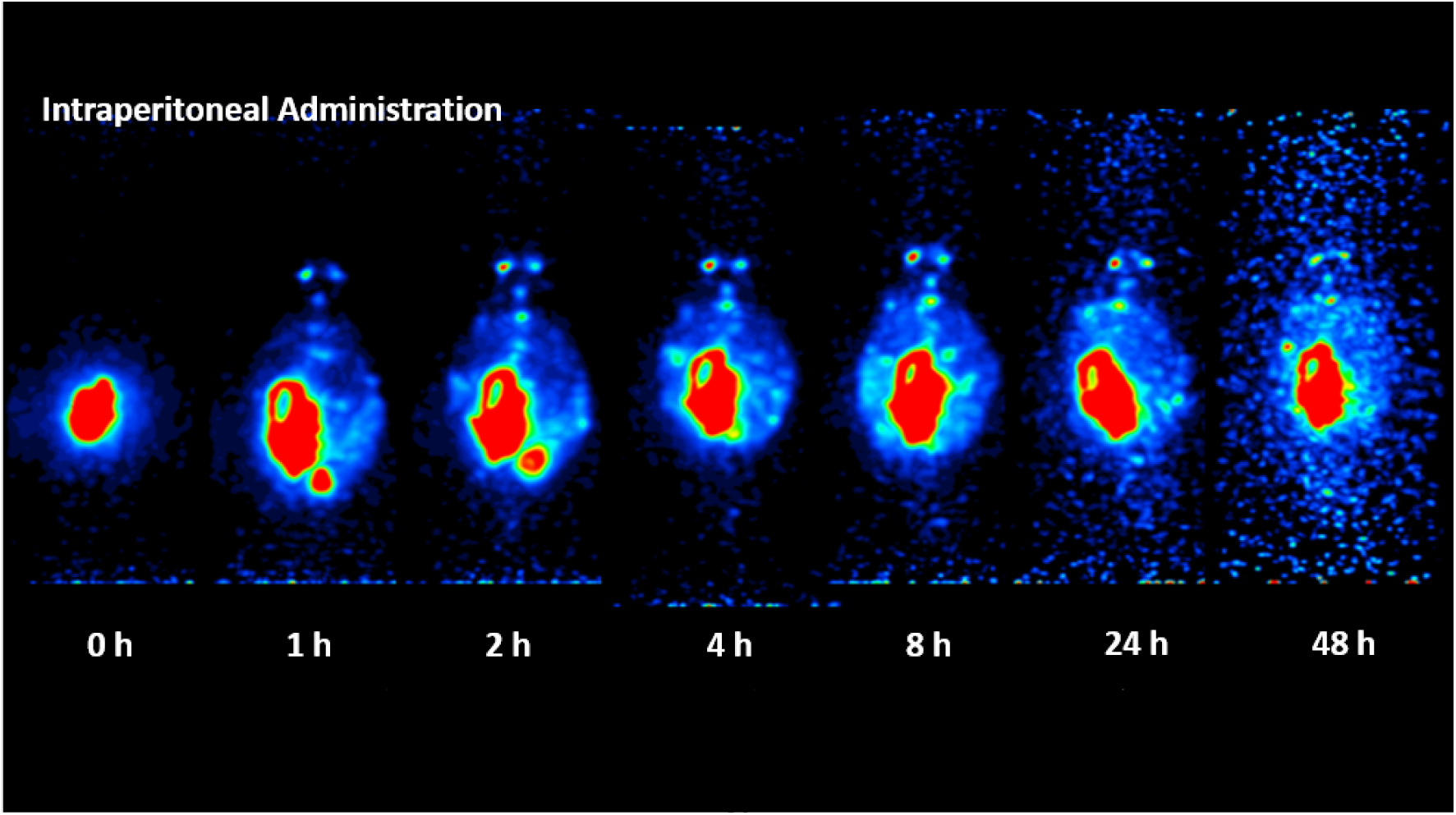
Biodistribution of ^64^Cu labeled OMVs after intraperitoneal administration in mice.

**Supplementary Figure 4.**
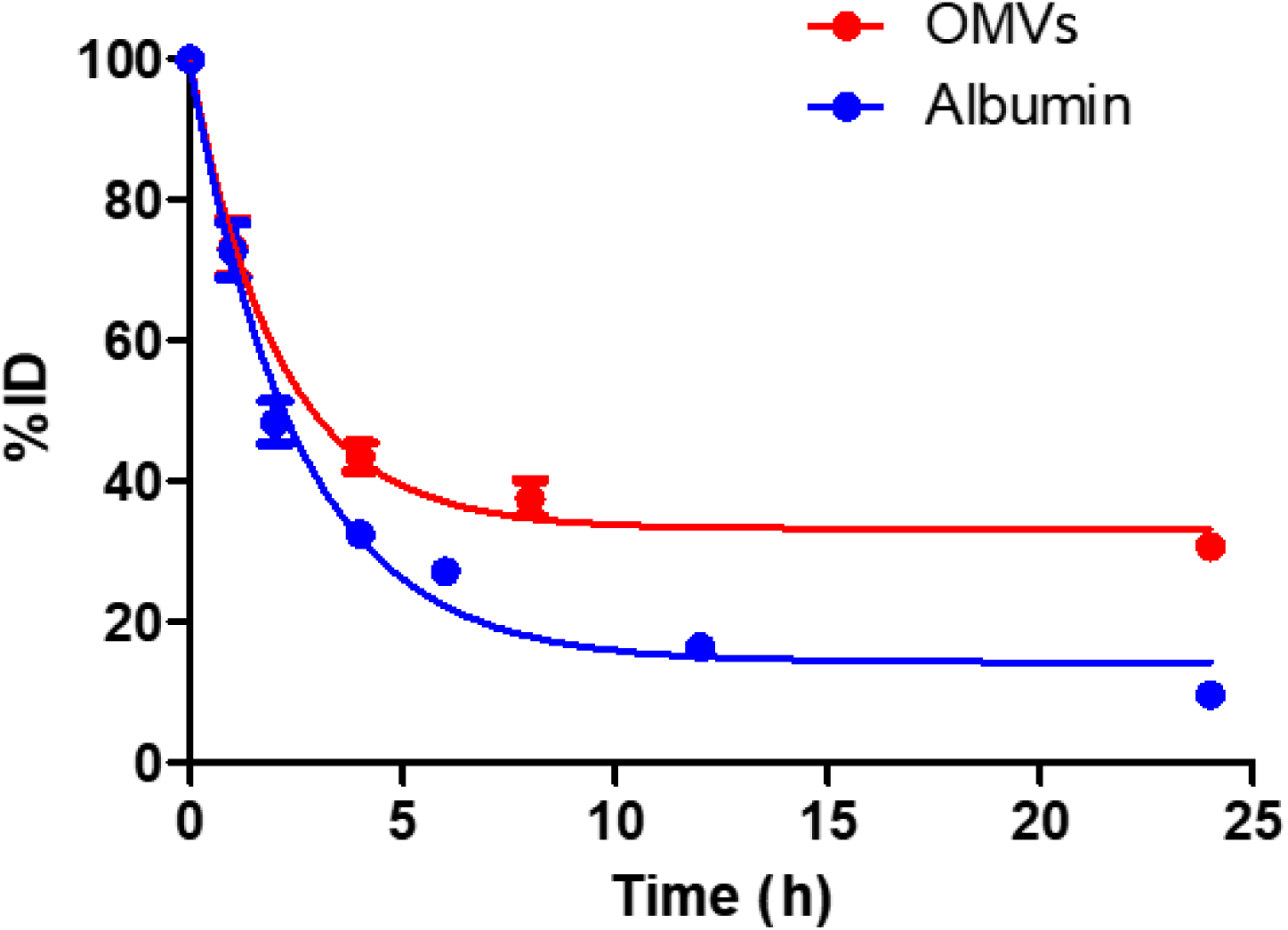
Kinetics of intra-CSF space radioactivity %ID over the injected amount or ^64^Cu labeled OMVs after intrathecal administration in mice (n=3). Data of kinetics of intra-CSF space radioactivity of ^64^Cu labeled albumin were derived from our results reported in Scientific Reports (Sarker et al., 2023). Drainage of ^64^Cu labeled OMVs were slower than ^64^Cu labeled albumin.

**Supplementary Table 1.**
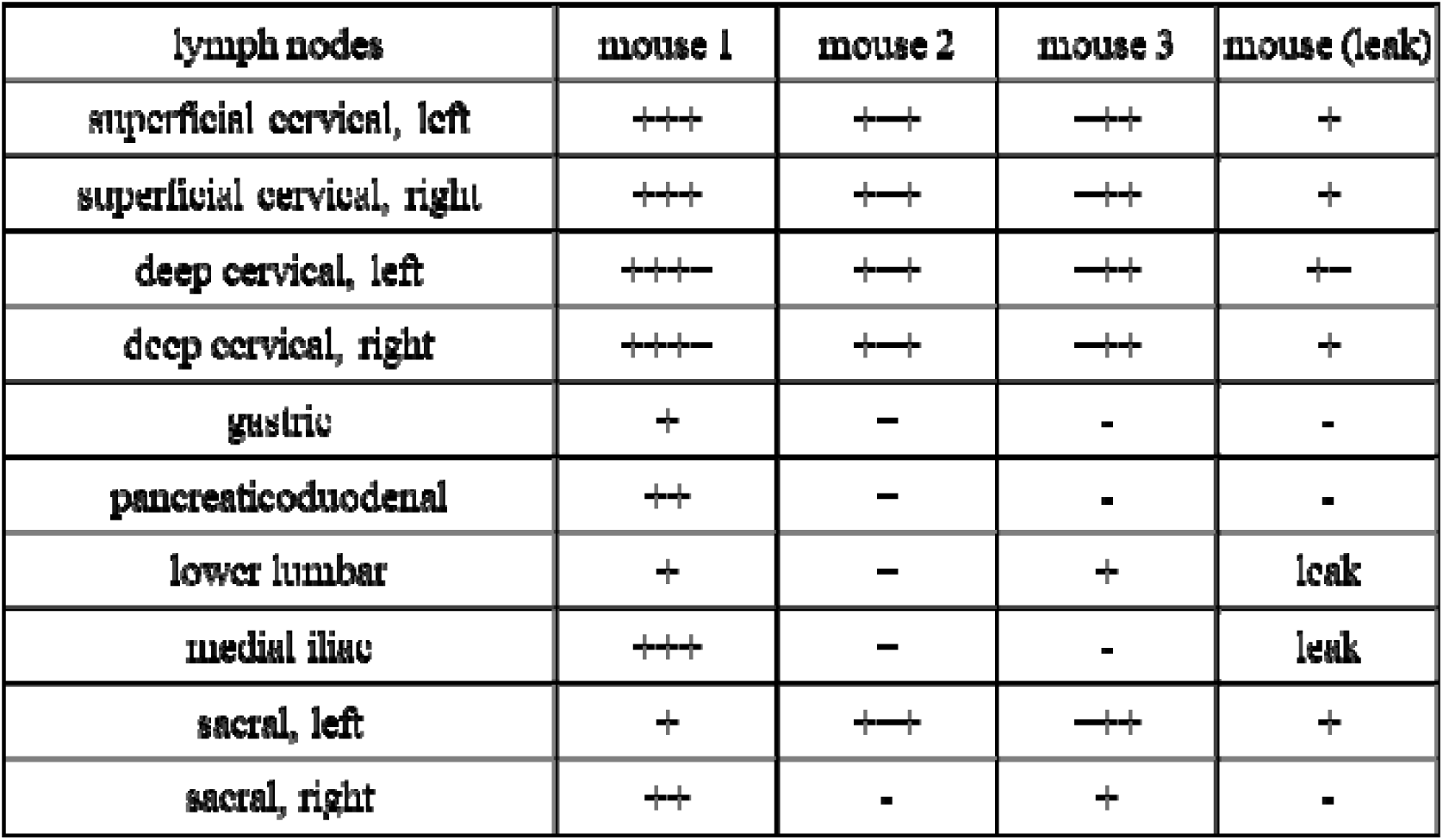
Ten lymph nodes and their uptake of ^64^Cu-labeled OMVs OMVs after intrathecal administration. Lumbar puncture was leaked in the fourth mouse (Suppl. Fig. 10). Intensity was graded semi-quantitatively by the first author, ranging from - to ++++. OMV: outer membrane vesicles.

