## Supplementary material for "Lymphatic drainage of cerebrospinal fluid using lymph node seeker ^64^Cu-labeled Gram-negative bacterial extracellular vesicles PET": Suppl Movies 1 to 10

### Slide 1
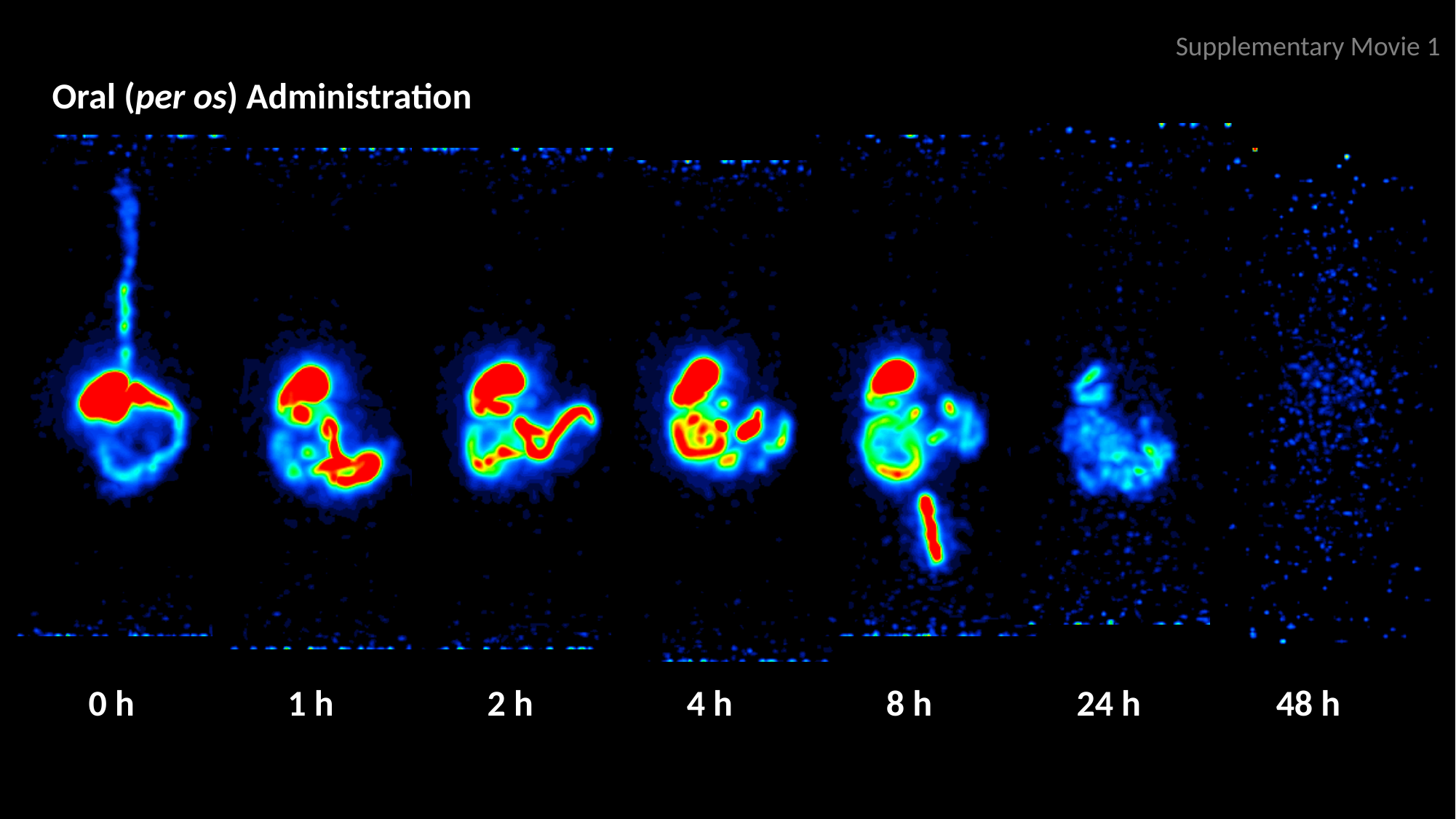

Supplementary Movie 1
Oral (per os) Administration
0 h
1 h
2 h
4 h
8 h
24 h
48 h

### Slide 2
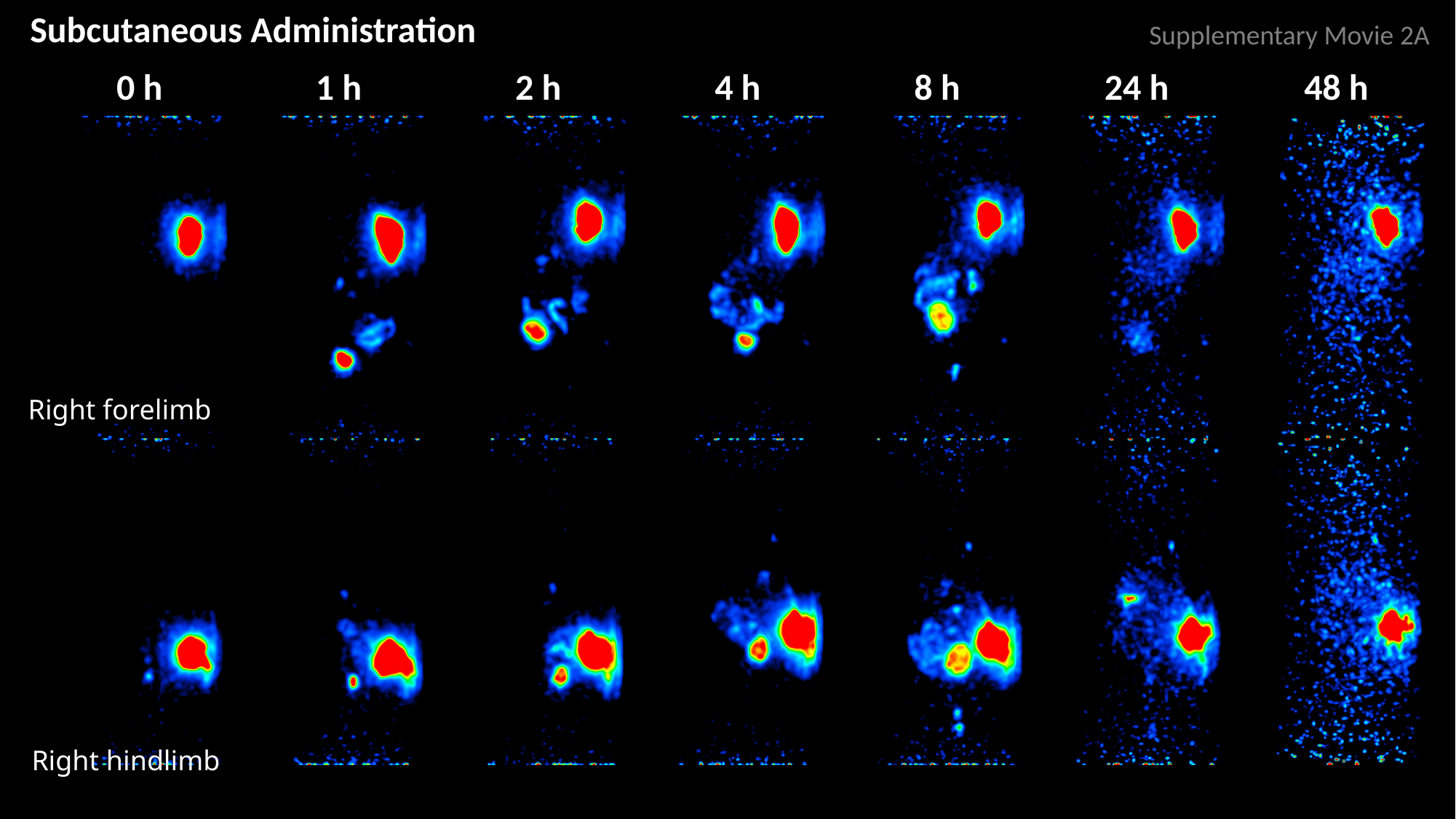

Subcutaneous Administration
Supplementary Movie 2A
0 h
1 h
2 h
4 h
8 h
24 h
48 h
Right forelimb
Right hindlimb

### Slide 3
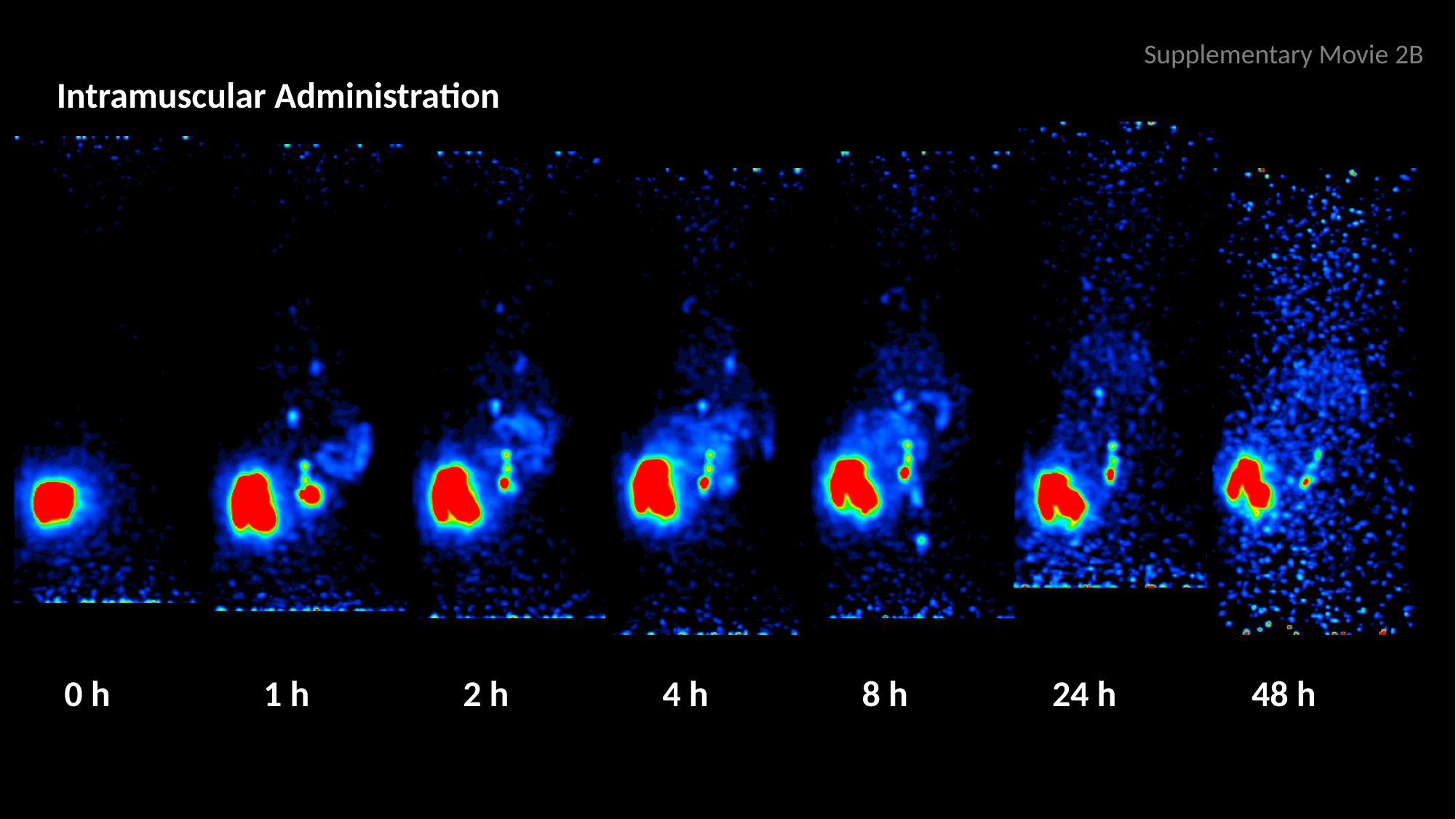

Supplementary Movie 2B
Intramuscular Administration
0 h
1 h
2 h
4 h
8 h
24 h
48 h

### Slide 4
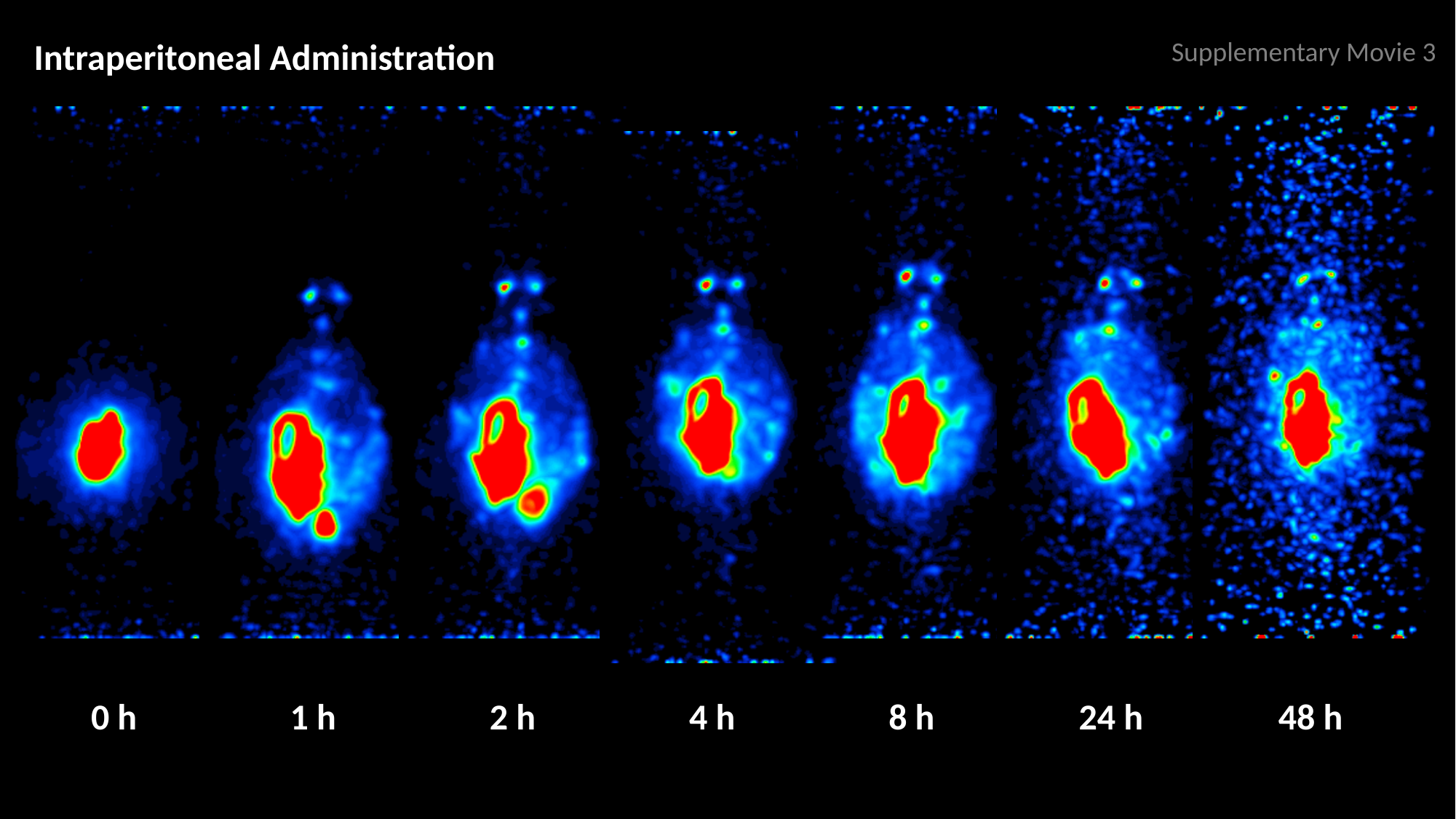

Intraperitoneal Administration
Supplementary Movie 3
0 h
1 h
2 h
4 h
8 h
24 h
48 h

### Slide 5
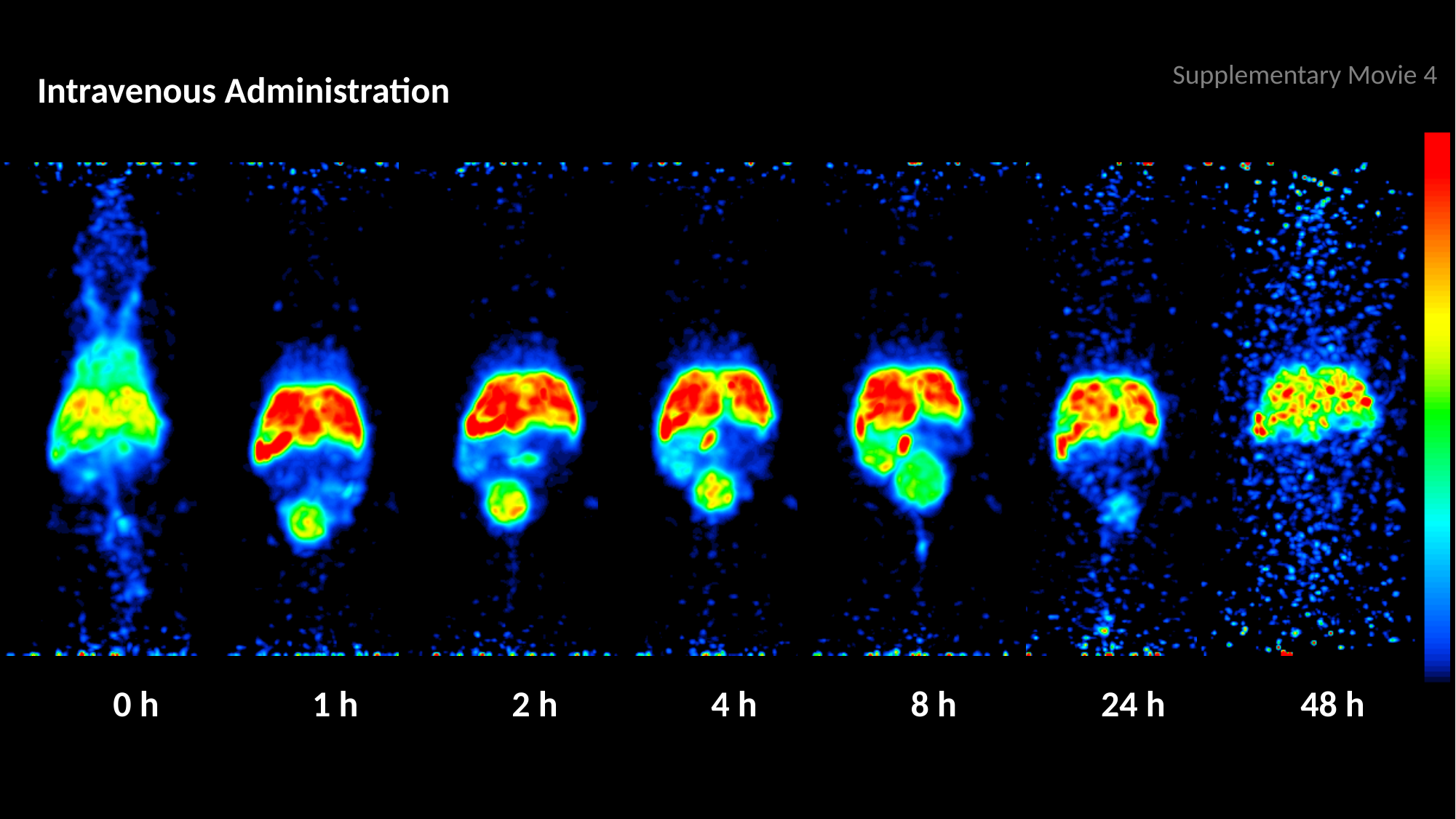

Supplementary Movie 4
Intravenous Administration
0 h
1 h
2 h
4 h
8 h
24 h
48 h

### Slide 6
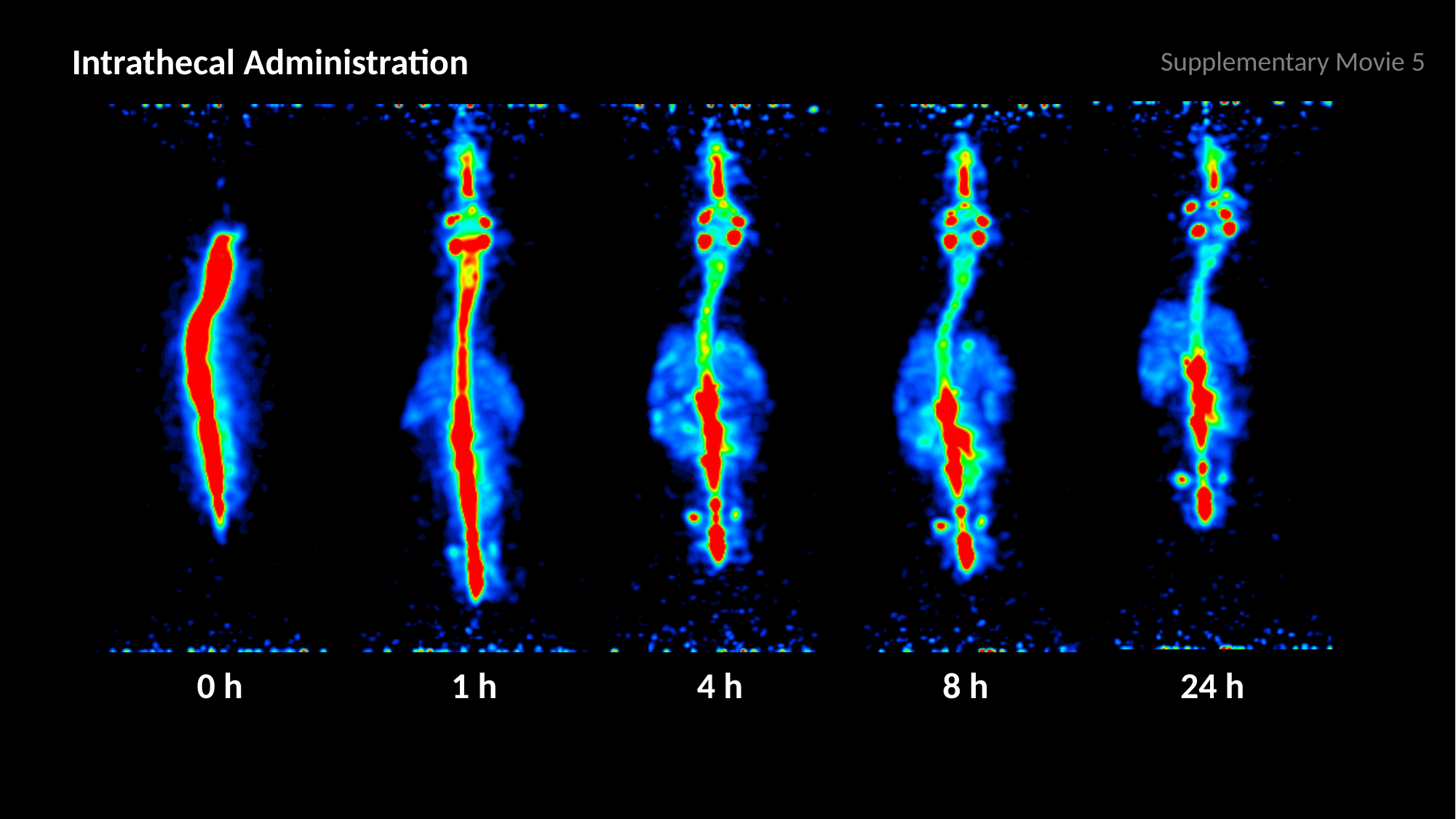

Intrathecal Administration
Supplementary Movie 5
0 h
1 h
4 h
8 h
24 h

### Slide 7
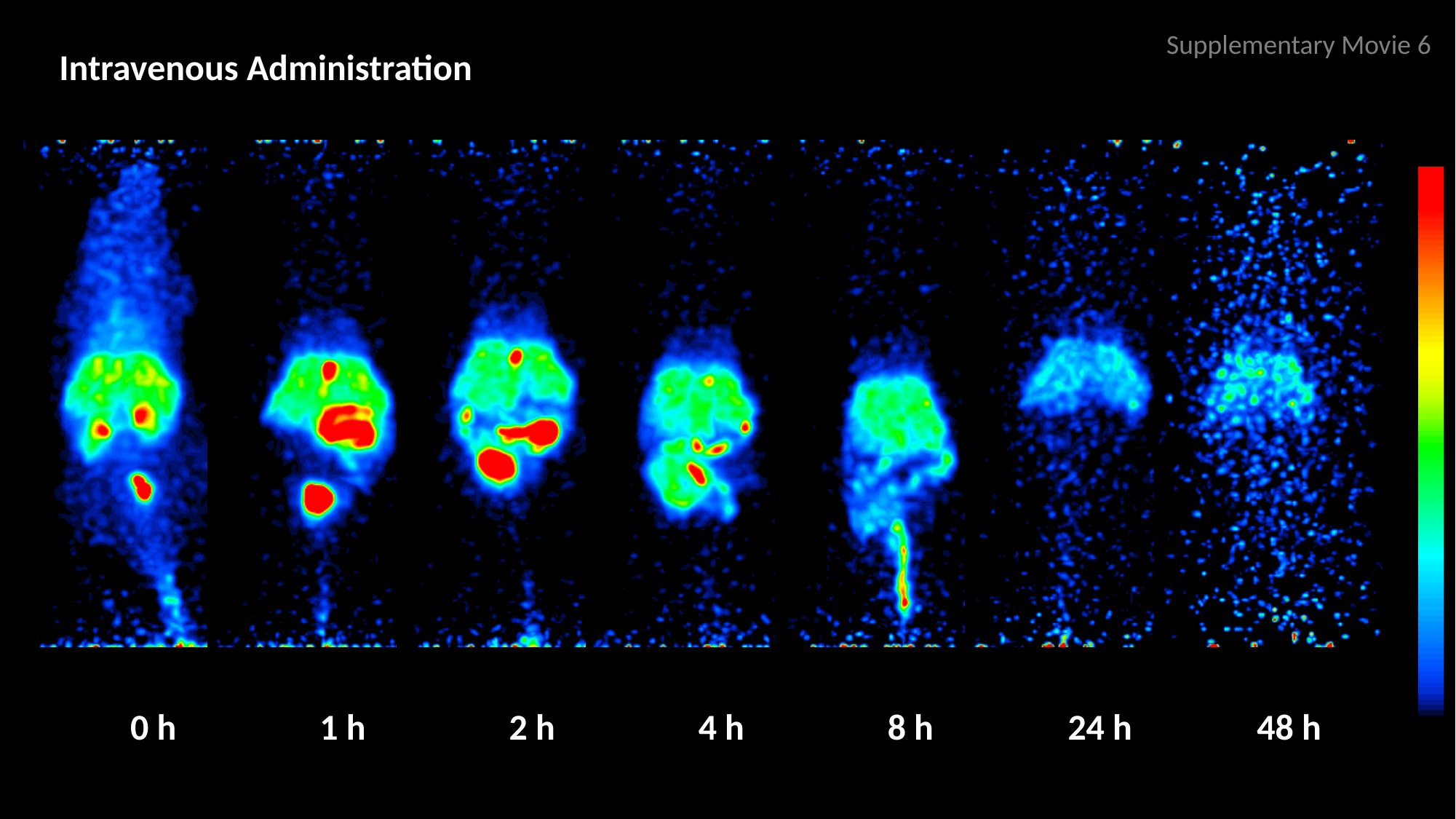

#
Supplementary Movie 6
Intravenous Administration
0 h
1 h
2 h
4 h
8 h
24 h
48 h

### Slide 8
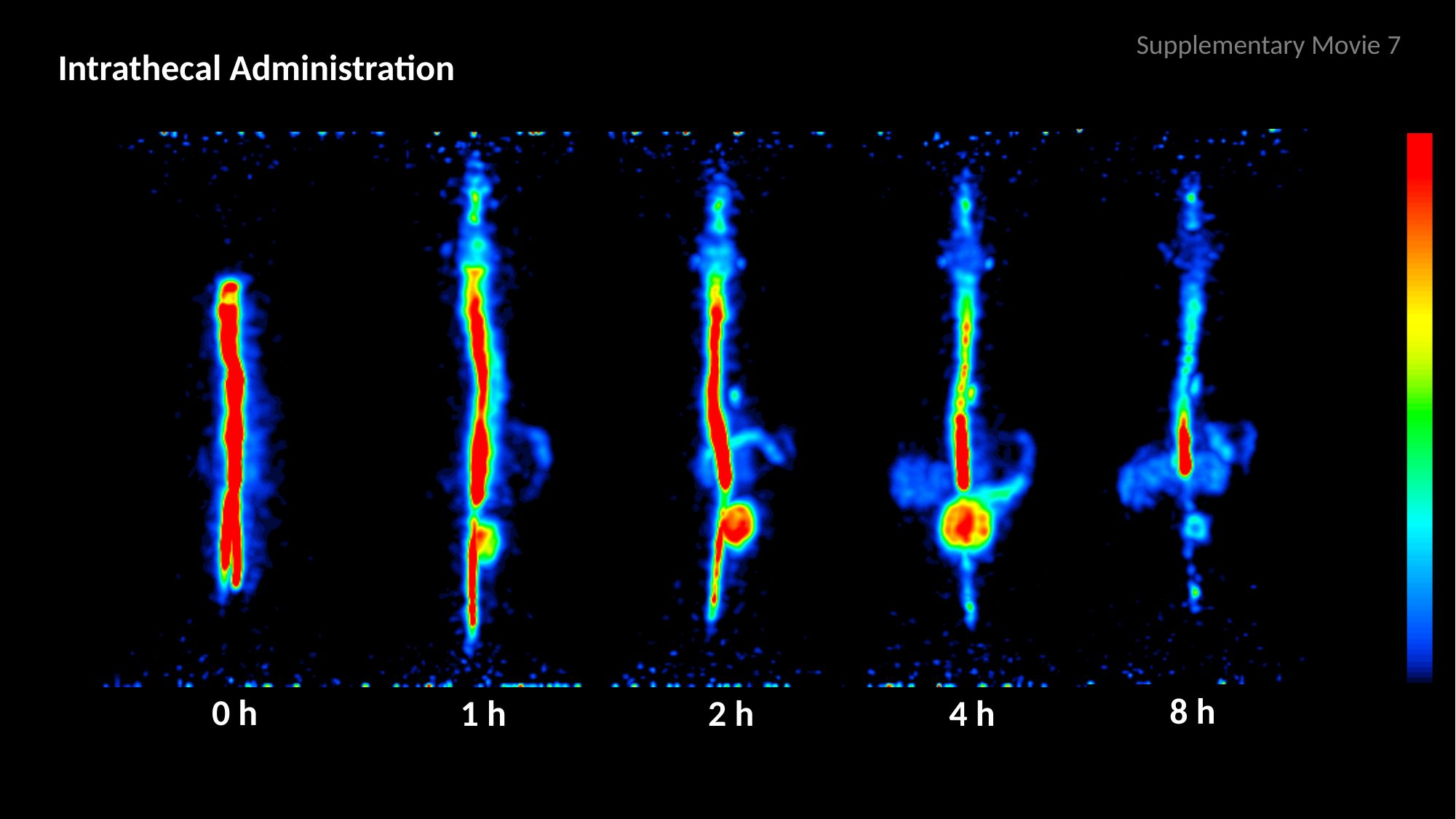

Supplementary Movie 7
Intrathecal Administration
8 h
0 h
1 h
2 h
4 h

### Slide 9
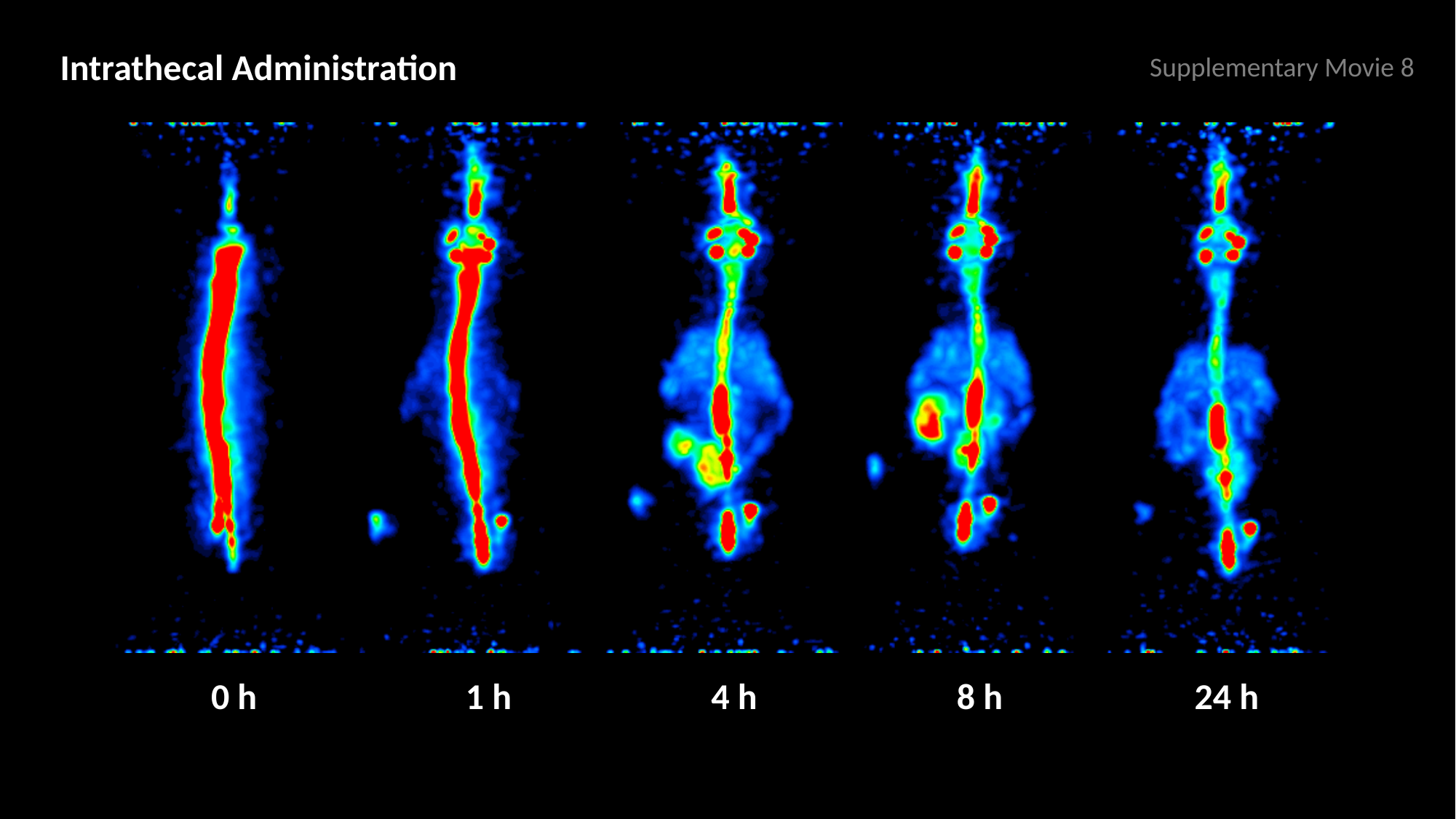

Intrathecal Administration
Supplementary Movie 8
0 h
1 h
4 h
8 h
24 h

### Slide 10
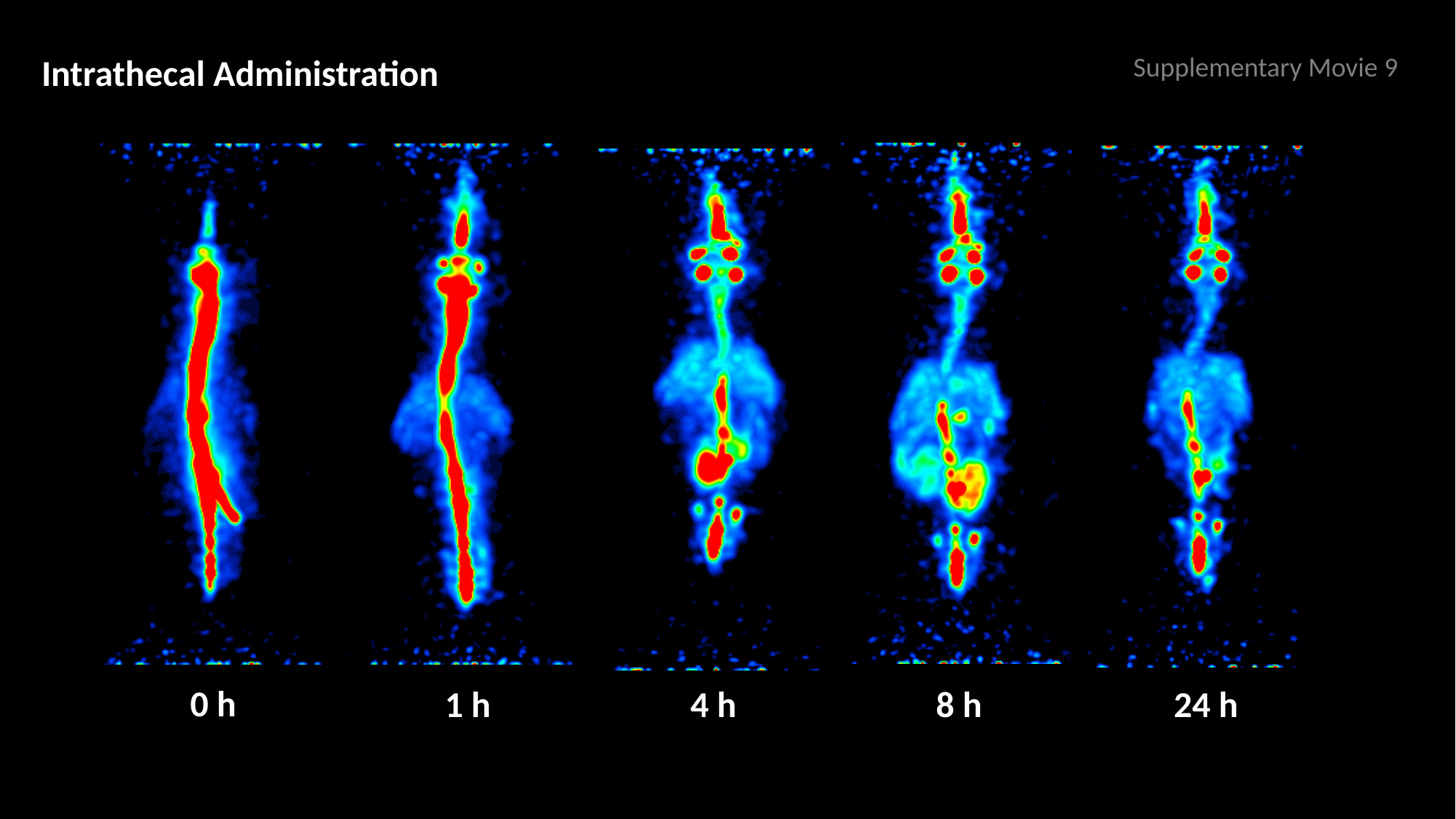

Supplementary Movie 9
Intrathecal Administration
0 h
1 h
4 h
8 h
24 h

### Slide 11
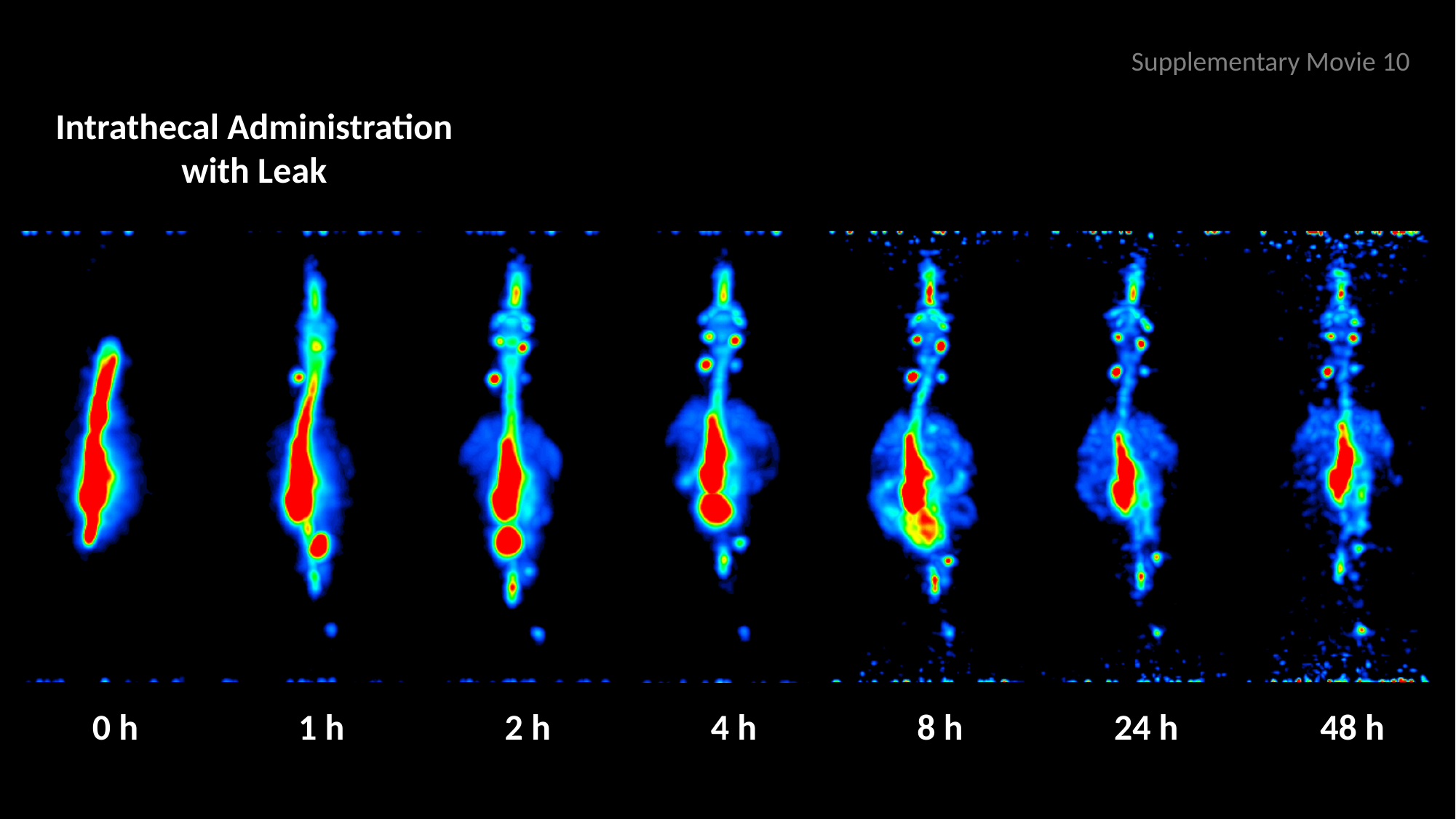

Supplementary Movie 10
Intrathecal Administration with Leak
0 h
1 h
2 h
4 h
8 h
24 h
48 h
